# Generation of human iPSC-derived motor neurons with defined regional identities across the rostrocaudal axis

**DOI:** 10.64898/2026.10.27.761477

**Authors:** Ken Ning, Ian Tindel, Brijesh Kumar Singh, Benisa Zogu, Ritchie Ho

**Affiliations:** Board of Governors Regenerative Medicine Institute, Cedars Sinai Health Sciences University, Los Angeles, CA 90048; Center for Neural Science and Medicine, Cedars Sinai Health Sciences University, Los Angeles, CA 90048; Department of Biomedical Sciences, Cedars Sinai Health Sciences University, Los Angeles, CA 90048; Department of Neurology, Cedars Sinai Health Sciences University, Los Angeles, CA 90048

## Abstract

This study establishes a unified pipeline for generating human induced pluripotent stem cell (iPSC)-derived motor neurons (iMNs) with defined hindbrain, cervical, and lumbar identities to support anatomically resolved studies of motor neuron development and degeneration. Conventional differentiation methods can efficiently produce iMNs but provide limited control over rostrocaudal identity and frequently generate populations with heterogeneous HOX profiles. We addressed this limitation by optimizing developmental morphogen combinations and treatment windows, establishing three complementary protocols that reproducibly generated hindbrain/rhombomere-associated motor neurons (RiMNs), cervical motor neurons (CiMNs), and lumbar motor neurons (LiMNs) across iPSC lines from six non-diseased donors balanced by sex. Transcriptomic and protein analyses demonstrated graded rostrocaudal specification from RiMNs through CiMNs to LiMNs, together with region-appropriate motor neuron subtype features. These identities showed the expected relationships to human week 9 fetal rostrocaudal HOX programs. High-density multielectrode array analysis further revealed synchronized network firing, electrically active neurite networks, and enhanced axonal signal propagation in the regional iMN populations compared with conventionally differentiated iMNs. Spike-sorted recordings additionally revealed protocol-associated differences in single-neuron electrophysiological profiles. Together, this three-protocol pipeline enables the reproducible generation and comparative analysis of anatomically defined human motor neuron populations and provides region-matched cellular models for investigating motor neuron development, selective degeneration, and therapeutic responses.

## Introduction

Motor neurons (MNs) acquire distinct rostrocaudal identities along the hindbrain and spinal cord that shape their subtype specification, peripheral targets, and functional properties^1^. These identities are encoded in part by spatially and temporally coordinated HOX gene expression and interact with transcriptional programs that specify motor columns and connectivity^2,3^. Consequently, MNs innervating cranial, axial, forelimb, and hindlimb musculature differ in their developmental histories and molecular states^4,5^. Reproducing this diversity is important for studying human motor system development and modeling disorders characterized by region-selective MN vulnerability.

Human induced pluripotent stem cell (iPSC)-derived MNs provide an experimentally accessible platform for investigating MN biology across defined genetic backgrounds. Established protocols recapitulate neural induction, ventralization, and MN specification but frequently generate heterogeneous rostrocaudal identities^6^. Transcriptomic datasets suggest that many iPSC-derived MN preparations are biased toward hindbrain/rhombomere and cervical programs, with variable activation of posterior HOX genes and likewise variable silencing of anterior HOX genes^7–9^. This heterogeneity complicates comparisons across experiments, limits the generation of MN populations associated with defined anatomical levels and innervation targets, and may confound the identification of disease-associated signatures^10^.

During embryogenesis, rostrocaudal neural identity is established through the coordinated action of developmental morphogens^5^. Retinoic acid, WNT, fibroblast growth factor, and related pathways regulate HOX activation, while ventralizing signals promote MN specification within regionally patterned progenitors^11–15^. Posterior spinal identities can also arise through neuromesodermal progenitors before neural commitment^14–17^. These developmental principles provide a framework for regional patterning in vitro, but their implementation requires precise control of morphogen combinations and exposure windows^18^. Regional identity must also be validated beyond individual HOX markers by assessing coherent transcriptional programs, MN subtype features, and reproducibility across genetic backgrounds. Human in vivo spatial transcriptomic datasets provide anatomically resolved references for assessing whether patterned iMNs acquire appropriate rostrocaudal programs. Functional characterization is equally important because molecular specification alone does not establish whether these cells are competent to develop coordinated network activity or support axonal signal propagation.

Here, we developed morphogen-based protocols for generating human iPSC-derived MNs with hindbrain/rhombomere-associated (RiMN), cervical (CiMN), and lumbar (LiMN) identities and compared them with conventionally directed iPSC-derived MNs (diMNs)^9,10^. Across six genetically distinct, non-diseased, and sex-balanced iPSC lines^19^, bulk transcriptomic and protein analyses demonstrated graded rostrocaudal identities from RiMNs through CiMNs to LiMNs while preserving robust MN differentiation and promoting region-appropriate features of MN subtypes including lateral and medial motor columns. Existing regionalization methods have substantially advanced the understanding of human MN development and differentiation and have provided specialized models for anatomically defined functions and disease contexts^17,18,20–25^. For example, phrenic MN models with defined molecular and functional identities offer valuable systems for investigating the development of diaphragm-innervating MNs and their involvement in motor neuron degenerative diseases^26^. Nevertheless, previously reported approaches optimized for different objectives involve distinct experimental tradeoffs, including terminal spheroid or organoid formats, incompletely resolved rostrocaudal heterogeneity, or limited preservation of limb-innervating lateral motor column identity. The three protocols developed here complement these models by establishing reproducible hindbrain, cervical, and lumbar HOX programs, preserving region-appropriate MN subtype features, and limiting 3D culture to a transient patterning phase before maturation as adherent 2D monolayers. This endpoint format facilitates longitudinal monitoring of cellular morphology, electrophysiological recording, and axon tracking across connected neuronal networks, while supporting broader implementation across laboratories and downstream applications. Together, these findings establish a reproducible platform for generating molecularly and functionally characterized human iMNs with experimentally controlled rostrocaudal identities.

### Morphogen-based patterning generates iPSC-derived motor neurons with distinct rostrocaudal identities

MNs acquire distinct rostrocaudal identities during development that contribute to their anatomical localization, central and peripheral innervation targets, and functional specialization^1^ (Figure 1A). Analysis of the Answer ALS transcriptomic dataset^10,19^ comprising 508 diMN samples revealed broad HOX expression across hindbrain, cervical/brachial, thoracic, and lumbar programs; the majority of iMNs exhibited predominantly hindbrain/rhombomere and cervical identities and substantial heterogeneity in region-defining HOX genes (Figure 1B). These findings suggest that conventional differentiation approaches provide limited control over rostrocaudal MN specification.

**Figure 1.**
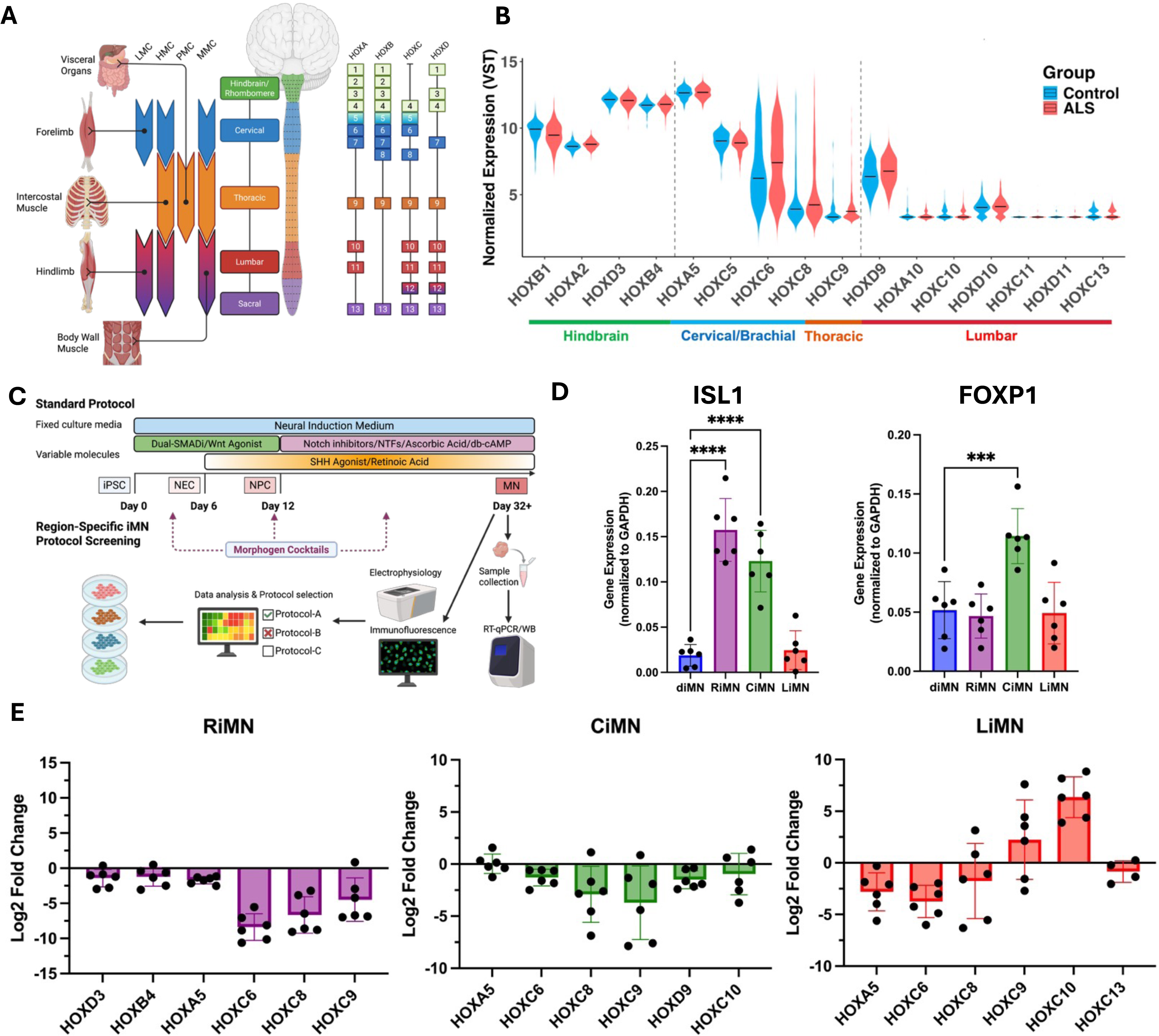
Morphogen-based patterning generates human iPSC-derived motor neurons with distinct rostrocaudal identities. **(A)** Schematic of motor neuron organization along the rostrocaudal axis, illustrating regional HOX expression, motor column identities, and peripheral innervation targets. **(B)** Expression distributions of selected HOX genes in conventionally differentiated iMNs from control and ALS donors in the Answer ALS dataset (158 control and 350 ALS samples). Violin plots show variance-stabilizing transformation (VST)-normalized expression, with genes arranged by their associated rostrocaudal domains. **(C)** Overview of the conventional iMN differentiation framework and the screening strategy used to establish region-specific protocols through optimization of morphogen combinations and treatment windows. **(D)** RT-qPCR analysis of *ISL1* and *FOXP1* expression in conventional diMNs and hindbrain/rhombomere-associated (RiMN), cervical (CiMN), and lumbar (LiMN) populations at day 32. Expression was normalized to *GAPDH* using the 2⁻ΔCt method. Statistical significance was assessed separately for each gene using one-way ANOVA followed by Dunnett’s multiple-comparisons test, comparing each regional iMN population with the diMN control. ***P < 0.001; ****P < 0.0001. **(E)** RT-qPCR analysis of selected HOX genes in RiMNs, CiMNs, and LiMNs at day 32. *GAPDH*-normalized expression is shown as log₂ fold change relative to the mean expression in diMNs. The horizontal line at zero indicates the diMN reference. For **(D)** and **(E)**, each point represents one independent iPSC line (n = 6 per group), and bars show mean ± SD.

We first characterized the temporal acquisition of neural, regional, and MN identities during conventional diMN differentiation. Pluripotency markers rapidly declined upon neural induction, accompanied by transient induction of neural stem and progenitor markers (Figure S1B). Rostral HOX genes emerged early, followed by cervical and more caudal HOX genes, consistent with progressive collinear HOX activation^12^ (Figures S1B and S1C). In parallel, MN markers including FOXP1, RALDH2, CHAT, and ISLET1 progressively increased during specification and maturation (Figure S1C). Transcriptomic profiling of selected diMN samples further confirmed robust neural and MN marker expression together with a broad HOX profile (Figure S1A). Thus, conventional diMN differentiation recapitulates progressive neural and MN specification but generates broad rostrocaudal identities.

To achieve greater control over regional identity, we introduced a transient 3D progenitor phase into the established iPSC-to-MN differentiation framework. As self-organized tissue architecture and mechanics can regulate developmental gene expression and modulate the response of human neural tissues to patterning cues^27,28^, we reasoned that transient aggregation might provide a tissue-like context in which rostrocaudal signals could be more effectively refined. We first tested whether this change in culture geometry was sufficient to alter MN specification by applying the conventional diMN media and morphogen regimen in a 3D-to-2D format (diMN 232D). Relative to adherent diMNs, diMN 232D cultures showed comparable ISL1 expression and retained rostral HOX expression, while exhibiting increased expression of selected cervical HOX genes, including HOXC6, HOXC8, and HOXC9 (Figure S1D). Thus, transient 3D culture alone did not establish a defined regional MN identity, but the partial shift in HOX expression suggested that the progenitor spheroid stage could provide a responsive developmental context for further optimization of rostrocaudal patterning.

We systematically optimized morphogen combinations, exposure windows, and basal medium components during transient 3D differentiation, leading to protocols for generating hindbrain/rhombomere-associated RiMNs and cervical CiMNs (Figure 1C). The protocols incorporated distinct region-specific patterning conditions, with LiMN differentiation proceeding through a neuromesodermal progenitor-like intermediate (Figure S1E). Following regional patterning, all three populations were plated as adherent monolayer cultures for MN maturation and downstream analyses. Across six iPSC lines from non-diseased donors, RiMN and CiMN spheroids expanded consistently while maintaining high roundness during the patterning period (Figures S1F and S1G, Table S4). LiMN spheroids similarly expanded during caudalization and resumed growth after mechanical chopping to control spheroid size^29–31^ (Figures S1H and S1I, Table S4).

We next assessed MN and regional identities generated by the optimized protocols. All three populations expressed ISL1, with significantly higher expression in RiMNs and CiMNs than in diMNs. CiMNs also showed increased FOXP1 expression, consistent with acquisition of limb-innervating lateral motor column features^32–34^ (Figure 1D). Analysis of rostrocaudal HOX expression revealed distinct regional profiles: RiMNs showed pronounced suppression of cervical and caudal HOX genes, including HOXC6, HOXC8, and HOXC9; CiMNs maintained cervical HOX expression while reducing more caudal HOXC8 and HOXC9; and LiMNs showed reduced rostral HOXA5 and HOXC6 together with increased caudal HOXC9 and HOXC10 (Figure 1E). These reciprocal HOX profiles demonstrate that controlled morphogen patterning enables the generation of iPSC-derived MNs with distinct hindbrain, cervical, and lumbar identities.

### Region-specific iMNs establish distinct rostrocaudal identities while maintaining motor neuron identity

To validate regional identity at the protein level, we examined HOXC6 and HOXC10 as markers of cervical and caudal HOX identities, respectively. Immunofluorescence revealed HOXC6 expression in diMNs and CiMNs, whereas HOXC6 was markedly reduced in RiMNs and LiMNs. HOXC10 staining was enriched in LiMNs (Figure 2A). Quantification showed a significantly higher proportion of HOXC6+ cells in CiMNs compared with diMNs and substantially fewer HOXC6+ cells in RiMNs and LiMNs (Figure 2B). LiMNs showed the highest proportion of HOXC10+ cells, significantly exceeding all other iMN populations (Figure 2C). HOXC10 mean fluorescence intensity was also significantly higher in LiMNs than in diMNs (Figure 2D). Bulk RNA-seq further showed reduced HOXC6 expression in RiMNs and LiMNs, maintained HOXC6 expression in CiMNs, and selective enrichment of HOXC10 in LiMNs, consistent with the RT-qPCR profiles in Figure 1E (Figures 2E and 2F).

**Figure 2.**
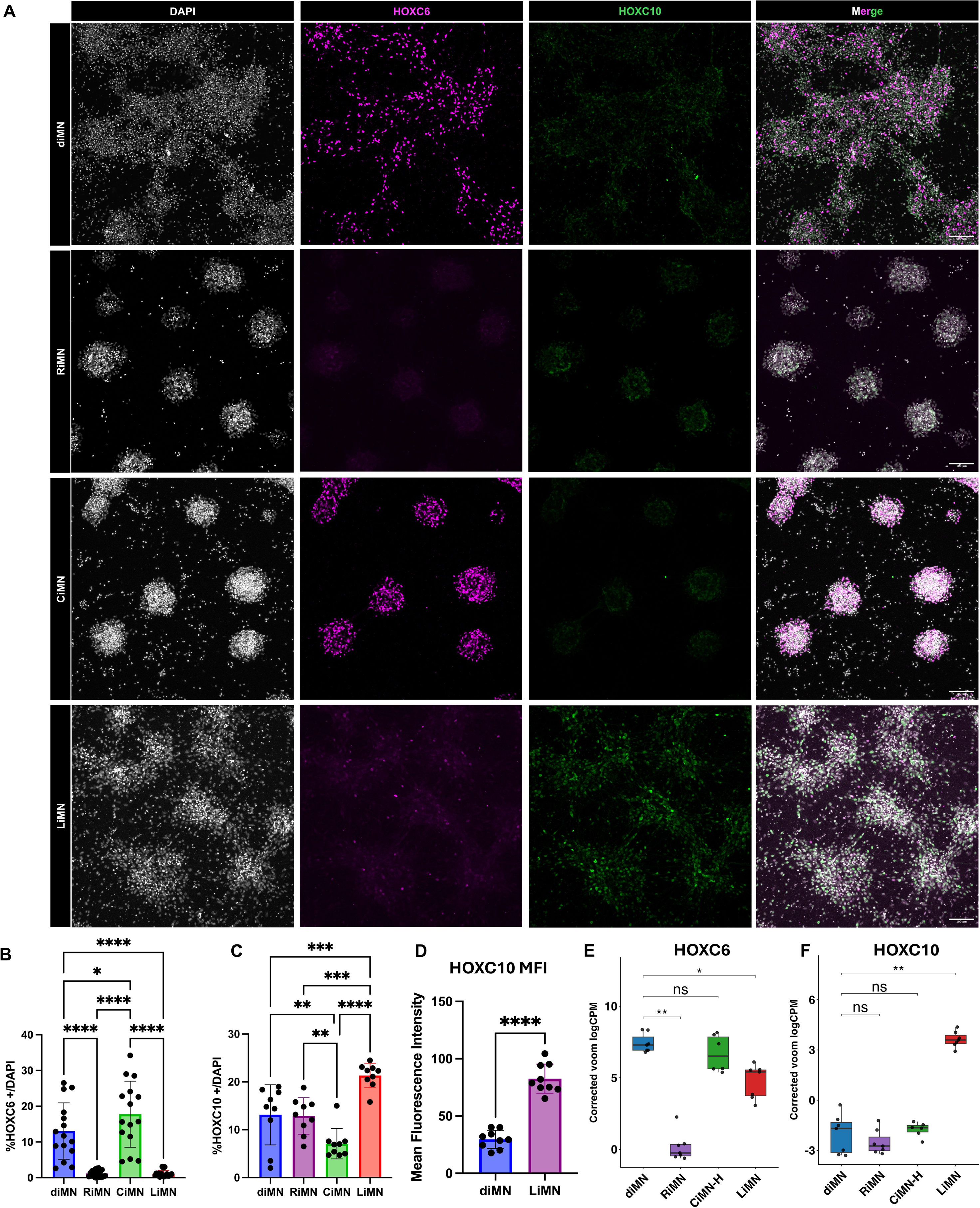
Regional iMN populations exhibit distinct HOXC6 and HOXC10 expression profiles. **(A)** Representative immunofluorescence images of conventional diMNs, RiMNs, CiMNs, and LiMNs at day 32, stained for HOXC6 (magenta) and HOXC10 (green), with DAPI counterstaining (white). Merged images are shown in the rightmost column. **(B and C)** Quantification of HOXC6-positive **(B)** and HOXC10-positive **(C)** cells as percentages of total DAPI-positive nuclei at day 32. Bars show mean ± SD. Statistical significance was assessed using one-way ANOVA followed by Tukey’s multiple-comparisons test. **(D)** Quantification of HOXC10 mean fluorescence intensity in diMNs and LiMNs at day 32. Statistical significance was assessed using a t-test. For **(B–D)**, each point represents one image, with three images analyzed per line across three independent iPSC lines per condition (nine images per condition). **(E and F)** Bulk RNA-seq expression of *HOXC6* **(E)** and *HOXC10* **(F)** at day 32 across the four iMN populations. Each point represents one independent iPSC line. Statistical significance was assessed using two-sided paired t-tests across matched cell lines, with Benjamini–Hochberg correction for multiple comparisons. *P < 0.05; **P < 0.01; ***P < 0.001; ****P < 0.0001; ns, not significant. Significance annotations in **(E)** and **(F)** indicate adjusted P-values. Scale bars in **(A)**, 100 µm.

To determine whether these selected markers reflected broader regional programs, we examined expression across the four HOX clusters. Sample-level heatmaps revealed reproducible protocol-associated HOX profiles across individual cell lines (Figures S2A and S2B). Group-mean profiles further demonstrated a graded rostrocaudal transition from RiMNs through CiMNs to LiMNs: RiMNs broadly suppressed posterior HOX genes, CiMNs maintained cervical expression while restricting more caudal programs, and LiMNs showed the strongest posteriorization, including increased HOXA9–HOXA11, HOXC10–HOXC11, and HOXD9–HOXD11 (Figure S2C). Together with the RT-qPCR and protein analyses, these data demonstrate that the protocols establish coordinated regional HOX programs rather than changes limited to individual markers.

Regional specification was also accompanied by MN subtype features appropriate to the corresponding rostrocaudal levels. CiMNs showed increased FOXP1 and ALDH1A2 expression, maintained LHX1, and reduced LHX3 relative to diMNs (Figures S2D–S2G), supporting enhanced specification of cervical lateral motor column-like MNs^35^. FOXP1 enrichment was consistent with the RT-qPCR results in Figure 1D and was confirmed by immunofluorescence, with CiMNs showing the strongest FOXP1 expression while LiMNs maintained expression comparable to conventional diMNs (Figure S2H). RiMNs showed reduced LHX1 and LHX3 expression and increased ISL2 and PHOX2B expression relative to diMNs, with PHOX2B enrichment further confirmed by immunofluorescence (Figures S2I and S2J). Together with suppression of posterior HOX programs, these features support acquisition of a hindbrain/rhombomere-associated MN identity^3,36^.

We next assessed whether regional patterning preserved MN differentiation and neurite development. All four populations expressed ISL1 and NEFH and formed extensive NEFH+ neurite networks (Figure S3A). Bulk RNA-seq showed increased ISL1 and NEFH expression in RiMNs and CiMNs relative to diMNs, with ISL1 expression consistent with the RT-qPCR results in Figure 1D (Figure S3B). Quantitative immunofluorescence further demonstrated higher proportions of ISL1+ cells and greater ISL1 signal intensity in RiMNs and CiMNs, whereas LiMNs remained comparable to diMNs (Figures S3C–S3E).

Morphological analysis of NEFH+ processes revealed significantly greater skeleton length and more network junctions per DAPI+ nucleus in RiMNs and CiMNs than in diMNs, consistent with increased neurite-network elaboration and complexity (Figures S3F and S3G, Table S5). LiMNs also formed extensive NEFH+ networks and remained comparable to diMNs across these normalized measures. CiMNs exhibited significantly lower local angular dispersion, indicating greater local alignment of NEFH+ processes (Figure S3H, Table S5). Although not statistically significant, LiMNs showed trends toward higher local angular dispersion and greater radial alignment than diMNs (Figures S3H and S3I, Table S5). These trends suggest a more radially oriented but locally heterogeneous neurite architecture, consistent with the morphology observed by NEFH immunostaining (Figure S3A). Collectively, these findings demonstrate that the optimized protocols establish coordinated rostrocaudal and region-appropriate MN subtype identities while preserving robust MN differentiation.

### Transcriptomic profiling identifies broad differentiation state and region-specific programs across iMN populations

Bulk RNA-seq analysis showed that the optimized regional protocols generated transcriptionally distinct iMN populations while preserving a shared neuronal differentiation program. PCA of batch- and cell-line-corrected expression profiles separated iPSCs from differentiated iMNs along PC1, whereas PC2 more clearly captured regional differences among iMNs, most prominently separating LiMNs from the rostral populations (Figure 3A). GO analysis of negatively loading PC2 genes identified significant enrichment of extracellular matrix and morphogenetic processes, including elastic fiber assembly, peptide cross-linking, cartilage condensation, and cell aggregation. Positively loading PC2 genes were displayed by fold-enrichment ranking and the highest-ranked terms included G protein-coupled glutamate receptor signaling and cGMP biosynthesis (Figure 3B). In contrast, PC1 loading genes were enriched for synaptic signaling and dense-core granule transport in one direction, and cell-cycle and early developmental processes in the other, indicating that PC1 primarily reflected the transition from proliferative iPSCs toward postmitotic neuronal states (Figure S4A).

**Figure 3.**
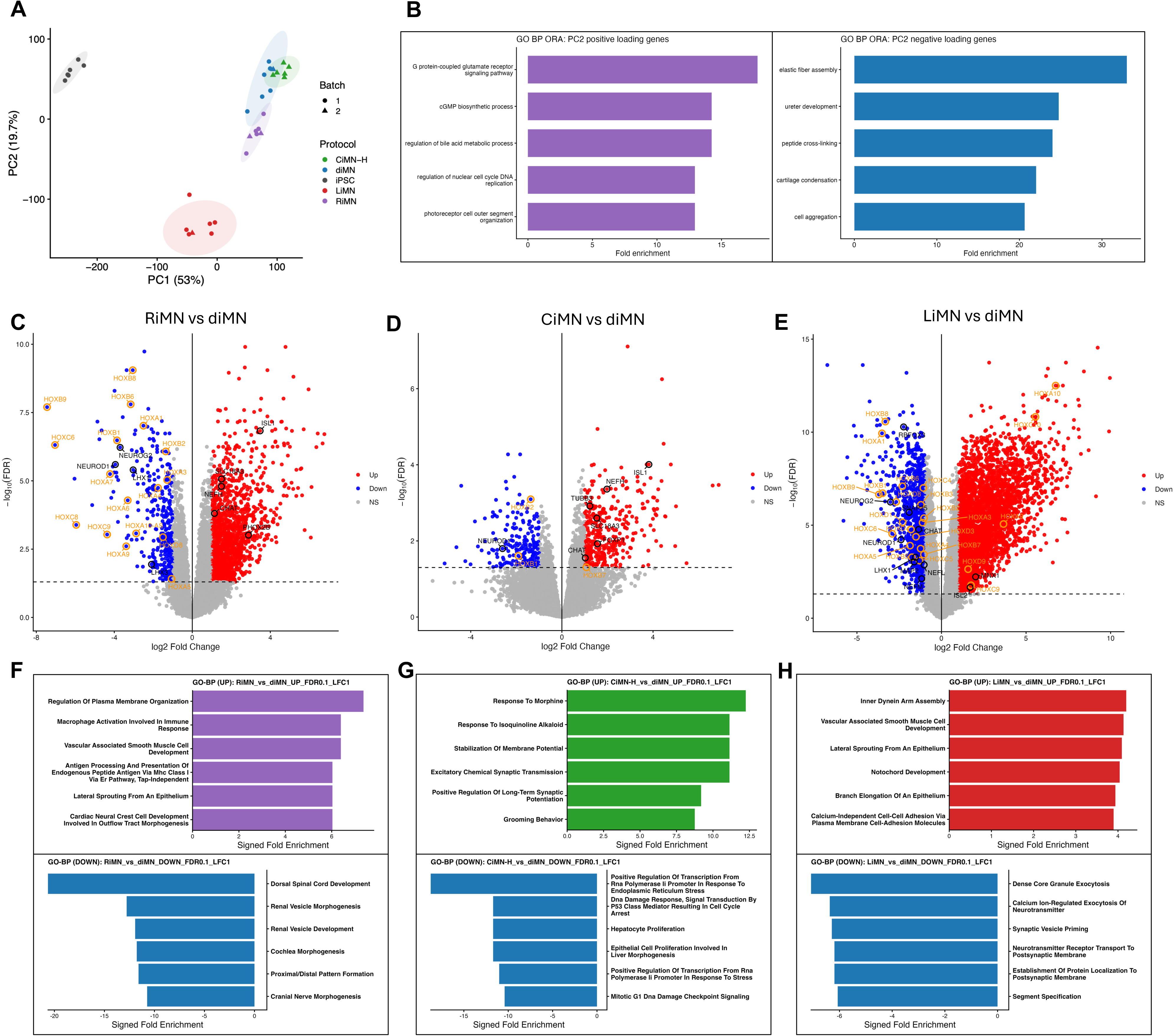
Transcriptomic profiling reveals distinct regional programs in iPSC-derived motor neurons. **(A)** Principal component analysis (PCA) of bulk RNA-seq profiles from iPSCs and day-32 diMNs, RiMNs, CiMNs, and LiMNs, using batch- and cell-line-corrected voom log₂ counts per million. Each point represents one sequenced sample. Colors indicate differentiation protocol, and symbols indicate sequencing batch. Shaded ellipses indicate 95% confidence regions. **(B)** Gene Ontology (GO) Biological Process over-representation analysis of the 250 genes with the strongest positive (left) and negative (right) PC2 loadings. The five highest-ranking terms by fold enrichment are displayed for each direction. **(C–E)** Volcano plots of differential gene expression in RiMNs **(C)**, CiMNs **(D)**, and LiMNs **(E)** relative to diMNs. Each point represents one gene, plotted by log₂ fold change and −log₁₀ false discovery rate (FDR). Red and blue indicate upregulated and downregulated genes, respectively, defined by FDR < 0.05 and absolute log₂ fold change > 1. Gray indicates genes not meeting both thresholds. **(F–H)** GO Biological Process over-representation analysis of genes upregulated (top) or downregulated (bottom) in RiMNs **(F)**, CiMNs **(G)**, and LiMNs **(H)** relative to diMNs. Input gene sets were defined using FDR < 0.1 and absolute log₂ fold change > 1. The six terms with the highest fold enrichment are displayed for each direction.

Heatmaps of the strongest PC1 and PC2 loading genes confirmed the separation of iPSCs from all iMN populations and highlighted the divergent transcriptomic profile of LiMNs (Figures S4B and S4C). Because diMNs, RiMNs, and CiMNs clustered closely in the full PCA, likely reflecting their shared differentiation framework, we repeated PCA after excluding iPSCs and LiMNs. In this restricted analysis, PC1 best distinguished RiMNs from CiMNs, with diMNs showing variation along this axis, and PC2 separated diMNs from RiMNs and CiMNs (Figure S4D). GO analysis of PC2 loading genes identified enrichment for developmental morphogenesis and regional patterning among positively loading genes, and for sensory perception, neuropeptide signaling, and circulatory processes among negatively loading genes (Figure S4E), indicating that RiMN and CiMN optimization produces measurable transcriptional divergence from conventional diMNs despite their shared protocol backbone.

Differential expression analysis further showed that each regional protocol remodeled the diMN transcriptome in a distinct manner. RiMNs showed broad downregulation of cervical and caudal HOX genes, including HOXC6, HOXC8, HOXC9, HOXB8, and HOXB9, consistent with suppression of posterior identity observed by RT-qPCR and immunostaining (Figures 1E, 2A, 2B, 2E, and 3C, Table S10). CiMNs exhibited a more selective transcriptional shift, consistent with the substantial cervical component already present in conventional diMNs, and showed increased expression of MN and subtype-associated genes, including ISL1, FOXP1, and ALDH1A2. Selected posterior HOX genes showed lower expression, but these decreases were not significantly differentially expressed (Figure 3D, Table S11). LiMNs displayed the strongest divergence from diMNs, with extensive upregulation of caudal HOX genes, including HOXD9, HOXC10, and HOXA10, and downregulation of rostral/cervical identity genes (Figure 3E, Table S12), consistent with HOXC10 protein enrichment and their distinct caudalizing differentiation trajectory. GO Biological Process enrichment analysis identified distinct patterns of developmental and neuronal gene expression across the regional protocols. Among the terms ranked highest by fold enrichment, RiMN-upregulated genes were associated with plasma membrane organization, vascular-associated smooth muscle cell development, and immune-related processes. RiMN-downregulated genes were enriched for dorsal spinal cord development, cochlea morphogenesis, and other developmental patterning processes (Figure 3F). CiMN-upregulated genes were enriched for stabilization of membrane potential, excitatory chemical synaptic transmission, and positive regulation of long-term synaptic potentiation, whereas downregulated genes were associated with DNA damage responses, stress-responsive transcription, and cell-cycle checkpoint signaling (Figure 3G). LiMN-upregulated genes were enriched for notochord development, epithelial morphogenesis, cell adhesion, and inner dynein arm assembly, whereas downregulated genes were associated with neurotransmitter release, synaptic vesicle priming, postsynaptic receptor trafficking, and segment specification (Figure 3H). Together, these enrichment patterns indicate protocol-associated differences in developmental, signaling, and synaptic gene programs. The diverse tissue and functional annotations reflect overlapping gene sets and do not, by themselves, establish corresponding cell identities or functional activities.

Pairwise comparisons among region-specific iMNs confirmed that LiMNs were the most transcriptionally distinct population, whereas RiMNs and CiMNs showed fewer differences from each other (Figures S4F–S4H). DEG-overlap analysis showed that each regional protocol produced both shared and protocol-specific changes relative to diMNs, with LiMNs containing the largest unique DEG set and RiMNs and CiMNs showing smaller but separable signatures (Figures S4I–S4K). Regional DEG sets showed substantial overlap with genes altered during conventional iPSC-to-diMN differentiation, alongside shared and protocol-specific changes relative to diMNs. These overlaps indicate that regional patterning modifies genes also involved in neuronal differentiation, although shared gene membership does not establish identical directions or magnitudes of expression change. Together, these bulk RNA-seq analyses extend mRNA and protein-level validation (Figures 1 and 2) by showing that morphogen-based patterning established distinct regional HOX, developmental, and neuronal subtype-associated transcriptomic states.

### Regional iMNs show HOX programs aligned with human fetal rostrocaudal reference profiles

To benchmark regional iMNs against human rostrocaudal development, we analyzed the Lawrence et al. week 9 fetal spatial transcriptomic dataset^37^. Spatial clusters were annotated using markers reported in the original study, identifying major neural and non-neural domains (Figures 4A and 4B). Cervical, thoracic, and lumbar regions contained broadly similar spatial-cluster classes but differed in their relative composition (Figures 4C and 4D; Figure S5A). ISL1- and ISL2-expressing profiles localized to a defined region of the fetal UMAP, identifying a transcriptionally coherent population enriched for putative MNs (Figure 4E). Expression of complementary markers, including OLIG2, NKX6-1, MNX1, NEFH, LHX1, LHX3, FOXP1, ALDH1A2, and PHOX2B, further characterized the progenitor, neuronal, and MN subtype-associated features represented within this population (Figure S5B). Accordingly, profiles expressing at least one canonical MN marker, MNX1, ISL1, or ISL2, were selected from each anatomical region. Regional dot-plot analysis confirmed expression of a broader panel of MN progenitor, MN, and subtype markers within neuron/glia-enriched fetal spatial profiles (Figure S5C). CHAT was not detectably expressed in the week 9 samples and was therefore excluded from subsequent marker-based correlations.

**Figure 4.**
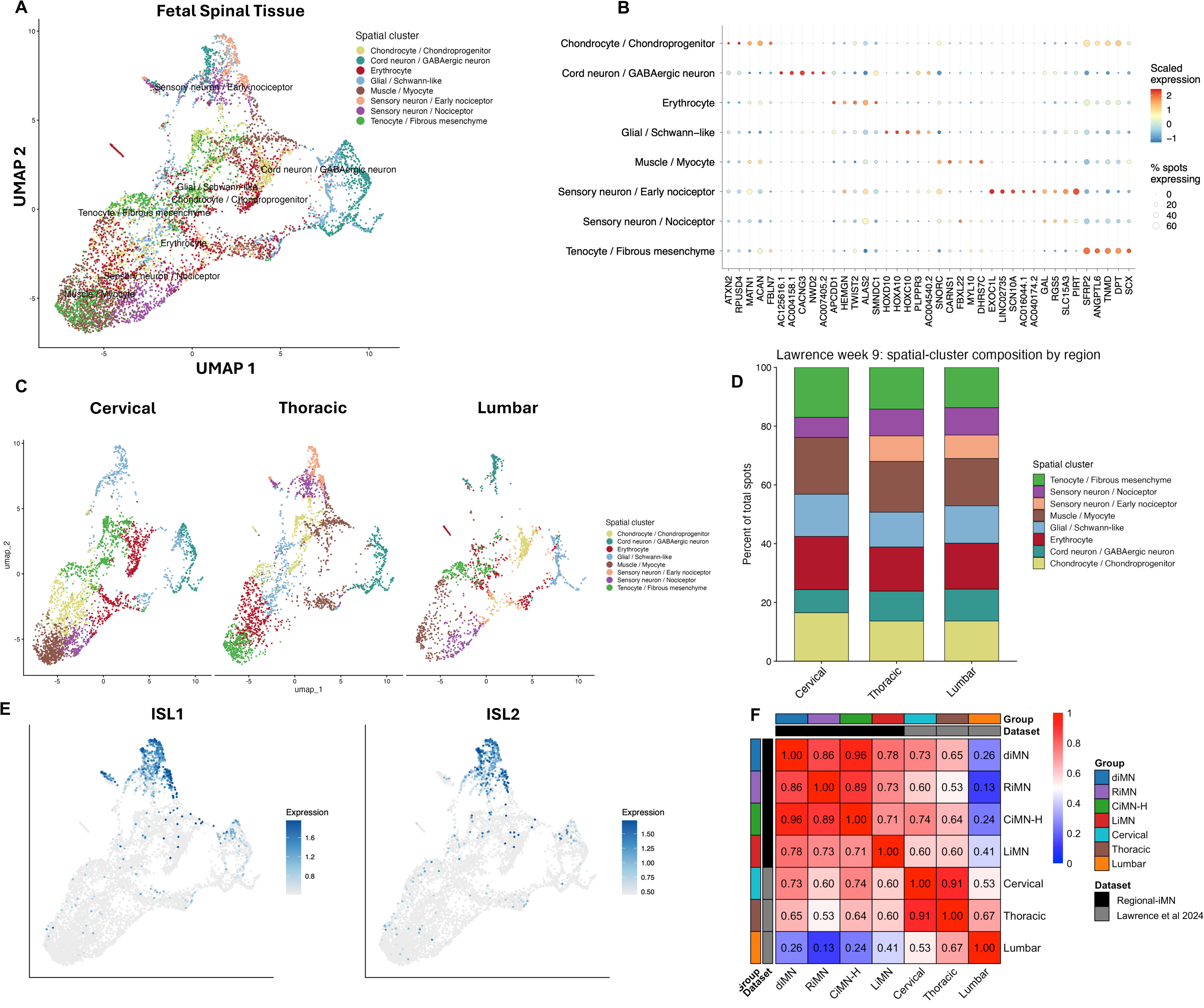
Regional iMNs show HOX expression profiles aligned with human fetal rostrocaudal identities. **(A)** UMAP of week-9 human fetal spinal cord spatial transcriptomic profiles from Lawrence et al. Each point represents one spatial spot, and colors indicate annotated spatial clusters. **(B)** Dot plot of representative marker-gene expression across annotated spatial clusters. Dot size indicates the percentage of spots expressing each gene, and color indicates scaled average expression. **(C)** UMAP of the annotated spatial clusters shown separately for cervical, thoracic, and lumbar regions. **(D)** Relative proportions of annotated spatial clusters across cervical, thoracic, and lumbar regions. **(E)** UMAP feature plots showing normalized expression of *ISL1* and *ISL2*. **(F)** Spearman correlation heatmap of HOX expression across day-32 diMNs, RiMNs, CiMNs, and LiMNs and week-9 fetal cervical, thoracic, and lumbar reference populations. Colors and values indicate Spearman correlation coefficients.

Regional HOX analysis of pseudobulk profiles aggregated from all week-9 fetal samples revealed distinct but overlapping cervical, thoracic, and lumbar programs, with progressive enrichment of posterior HOX genes toward the lumbar region (Figure S5D). These profiles indicate that human fetal rostrocaudal identities are graded rather than separated by completely discrete HOX boundaries at this developmental stage. All-gene Spearman correlations were uniformly high, reflecting broad transcriptional programs shared among MN populations and providing limited resolution of rostrocaudal identity (Figure S5E). Restricting the analysis to HOX genes more clearly resolved regional relationships: diMNs and CiMNs showed the greatest similarity to cervical fetal MNs, RiMNs showed reduced similarity to progressively posterior regions, and LiMNs displayed the highest lumbar correlation among the iMN protocols (Figure 4F). To determine whether inclusion of MN-lineage information altered these regional relationships, we repeated the correlation analysis using the shared, detectably expressed MN and MN progenitor markers (Figure S5C) together with HOX genes. The expanded analysis reproduced the overall structure of the HOX-only comparison: diMNs and CiMNs remained most similar to cervical profiles, RiMNs retained the lowest similarity to lumbar profiles, and LiMNs maintained the highest lumbar correlation among the iMN protocols (Figure S5F). Thus, although the additional markers confirmed shared MN-lineage features, they contributed limited additional rostrocaudal resolution. The preservation of the regional correlation pattern indicates that HOX programs remain the dominant source of positional information in this comparison, consistent with their central role in rostrocaudal specification during spinal MN development. Differences remaining between the in vitro and fetal profiles may reflect developmental timing, maturation state, and the spatial context of the fetal tissue. Together, these analyses show that morphogen-based patterning establishes graded regional programs in human iMNs that correspond to defined rostrocaudal identities in developing human MNs.

### HD-MEA profiling reveals enhanced network activity and axonal signal propagation in region-specific iMNs

To assess whether regional specification supports functional maturation, we profiled diMNs, RiMNs, CiMNs, and LiMNs using MaxWell high-density multielectrode array (HD-MEA) assays, enabling longitudinal measurements of spontaneous firing, network bursting, axonal propagation, neurite-associated activity, and spike-sorted single-neuron features (Figure 5A). Conventional diMNs showed limited spontaneous activity. In the representative diMN recording, the raster plot showed sparse firing events across electrodes, while the corresponding network firing-rate trace showed low activity without clear synchronized bursting (Figure 5B). Across six sex-balanced iPSC lines from non-diseased donors, diMNs maintained low mean firing rates and modest spike amplitudes over time (Figure 5C, Table S6), while axon tracking detected only limited propagating activity in selected recordings (Figures S7A and S7B, Table S6). This limited activity is consistent with previous reports that continuous 2D neuronal differentiation produces weaker electrophysiological phenotypes than spheroid-based culture^38–40^. By contrast, all three region-specific iMN populations developed synchronized spontaneous activity after monolayer plating. Representative raster plots showed dense, temporally aligned firing events across channels, and the corresponding network firing-rate traces showed rhythmic peaks consistent with coordinated population bursts (Figures 5D-F). Similar synchronized firing was observed in additional independent cell lines (Figures S6A-C), supporting reproducibility across genetic backgrounds. Longitudinal measurements showed increasing mean firing rates in RiMNs and CiMNs, while LiMNs maintained detectable synchronized activity with more moderate firing-rate changes (Figures 5J-5L, top; Tables S7–S9). Longitudinal spike-amplitude measurements showed temporal variation in RiMNs and CiMNs, whereas LiMN amplitudes remained comparatively stable (Figures 5J–5L, bottom; Tables S7–S9). Mean burst peak firing rates similarly increased during maturation in RiMNs and CiMNs (Figures S6G and S6H, top; Tables S7 and S8), whereas LiMNs showed smaller changes (Figure S6I, top; Table S9). Notably, all regional protocols included a transient spheroid phase before HD-MEA plating. This three-dimensional culture period aligns with prior studies showing that spheroid-based neural differentiation can enhance cell-cell interactions, early network organization, and activity-dependent maturation, thereby supporting synchronized firing and coordinated network bursting after plating^38,41–44^.

**Figure 5.**
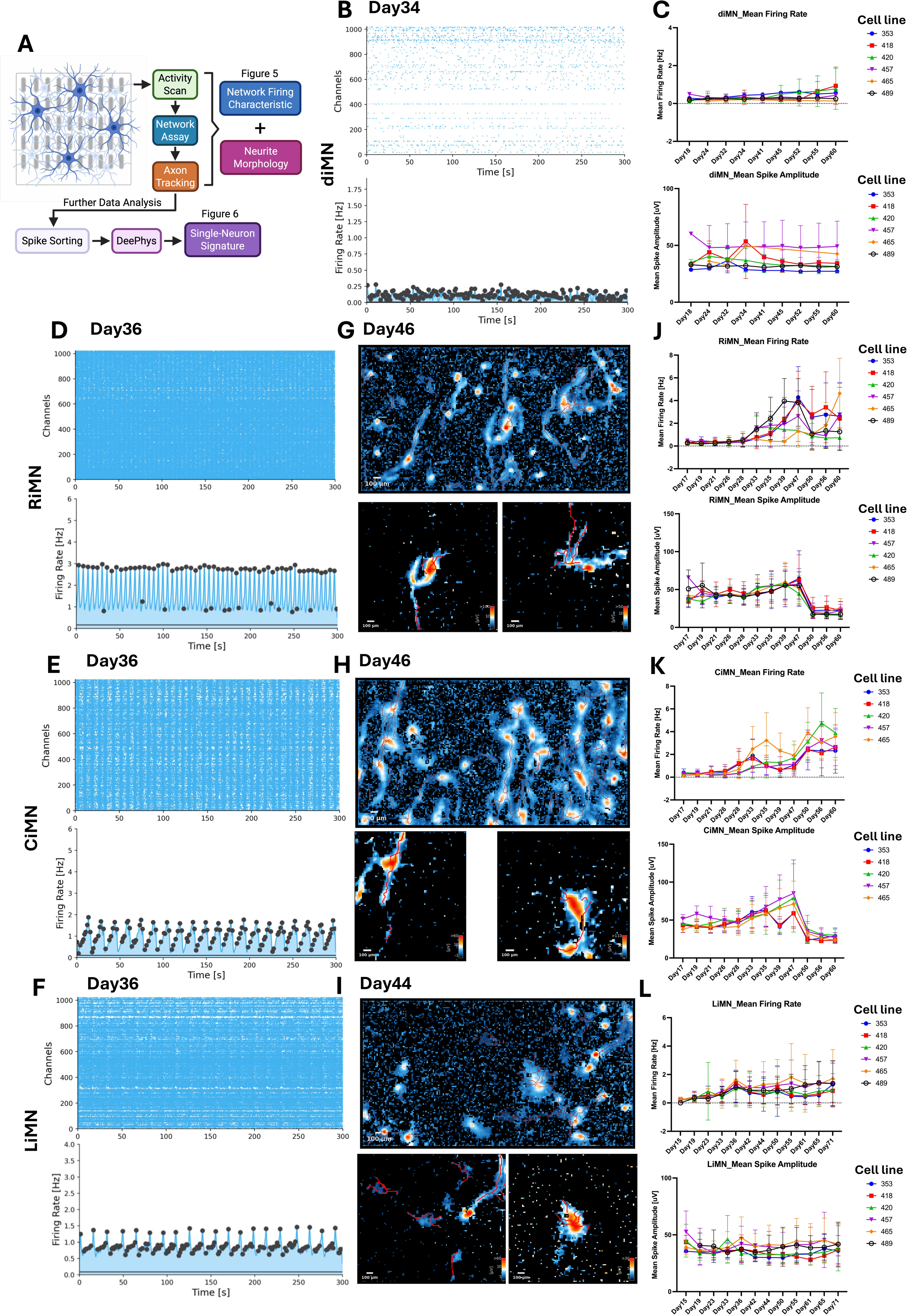
HD-MEA profiling reveals network activity and axonal signal propagation in regional iMNs. **(A)** Overview of the high-density multielectrode array (HD-MEA) workflow for assessing neuronal activity, network firing, axon tracking, and single-neuron electrophysiological features. **(B)** Representative spike raster plot (top) and corresponding network firing-rate trace (bottom) from diMNs at day 34. **(C)** Longitudinal measurements of mean firing rate (top) and mean spike amplitude (bottom) in diMNs. **(D–F)** Representative spike raster plots (top) and corresponding network firing-rate traces (bottom) from RiMNs (D), CiMNs (E), and LiMNs (F) at day 36. Raster plots show spike events across recording channels over 300 s. **(G–I)** Representative axon-tracking maps from RiMNs at day 46 (G), CiMNs at day 46 (H), and LiMNs at day 44 (I). Upper panels show full-field activity maps, and lower panels show examples of individual neuronal electrical footprints with tracked axonal trajectories. Scale bars, 100 µm. **(J–L)** Longitudinal measurements of mean firing rate (top) and mean spike amplitude (bottom) in RiMNs (J), CiMNs (K), and LiMNs (L). In (C, J–L), each colored trajectory represents one independent iPSC line (n = 5 for CiMNs; n = 6 for diMNs, RiMNs, and LiMNs). Data show mean ± SD, with differentiation days indicated on the x-axis.

Axon-tracking further showed that regional iMNs formed more extensive and electrically active neurite networks than diMNs. In the full-field axon-tracking maps, RiMNs and CiMNs displayed highly distributed activity along elongated and branched neurite-like trajectories, indicating complex electrically active networks on HD-MEAs (Figures 5G-H, top). These patterns were consistent with the more elaborate NEFH+ networks observed by immunostaining and the increased NEFH skeleton length and network junctions quantified in RiMNs and CiMNs (Figures S3A, S3F, and S3G, Table S5), demonstrating that their enhanced neurite morphology was accompanied by functional electrical propagation. LiMNs showed less extensive axonal activity than RiMNs and CiMNs but still exhibited detectable neurite-associated propagation across all tested lines, whereas diMNs showed limited and inconsistently detectable axonal activity (Figure 5I, top; Figures S6F and S7A, top, Table S6-9). Single-neuron firing event records further illustrated active axonal or neuritic paths, highlighting the length, branching, and propagation of firing-associated signals through these processes (Figures 5G-I, bottom; Figure S7A, bottom). Similar axon-tracking patterns in additional independent cell lines supported the reproducibility of enhanced neurite-associated activity in regional iMNs (Figures S6D-F, Table S6-9). These observations were consistent with longitudinal measurements showing increased total axon length in RiMNs and CiMNs but no consistent increase over time in LiMNs (Figures S6G–S6I, bottom; Tables S7–S9). Nevertheless, LiMNs exhibited greater total axon length than diMNs around day 45 (Figure S7D, middle; Tables S6 and S9).

Cross-protocol comparisons around day 45 confirmed broadly enhanced functional profiles in regional iMNs. RiMNs and CiMNs improved across nearly all HD-MEA readouts, including mean firing rate, burst peak firing rate, spike amplitude, conduction velocity, total axon length, and active area (Figures S7C and S7D, Table S6-8). These functional gains are consistent with the increased ISL1 expression and enhanced neurite-associated activity observed in RiMNs and CiMNs, and align with prior evidence that MN specification, neurite outgrowth, and network organization are associated with electrophysiological maturation^23,40,45–47^. LiMNs showed significant increases in conduction velocity, total axon length, and active area while maintaining firing-rate and spike-amplitude measures comparable to diMNs (Figures S7C and S7D, Table S6-9). Together, these data demonstrate that the regional differentiation protocols generate iMN cultures with synchronized network activity, enhanced neurite-associated electrical activity, and improved axonal signal propagation.

### Regional iMN cultures exhibit distinct single-neuron electrophysiological profiles

To compare single-neuron electrophysiological profiles among regional iMN populations, we analyzed spike-sorted HD-MEA recordings from RiMN, CiMN, and LiMN cultures at day 45 (Figure 5A). Comparison of the combined electrophysiological feature profiles revealed greater similarity among cultures generated using the same regional protocol than across protocols, supporting protocol-associated electrophysiological differences (Figure 6A, Table S13). Individual features nevertheless showed variation across cell lines and overlapping distributions among regional populations (Figures S8A–S8G). Summary feature profiles particularly distinguished LiMNs from RiMNs and CiMNs, with LiMNs exhibiting lower firing rates and longer mean interspike intervals (ISIs) (Figure 6B).

**Figure 6.**
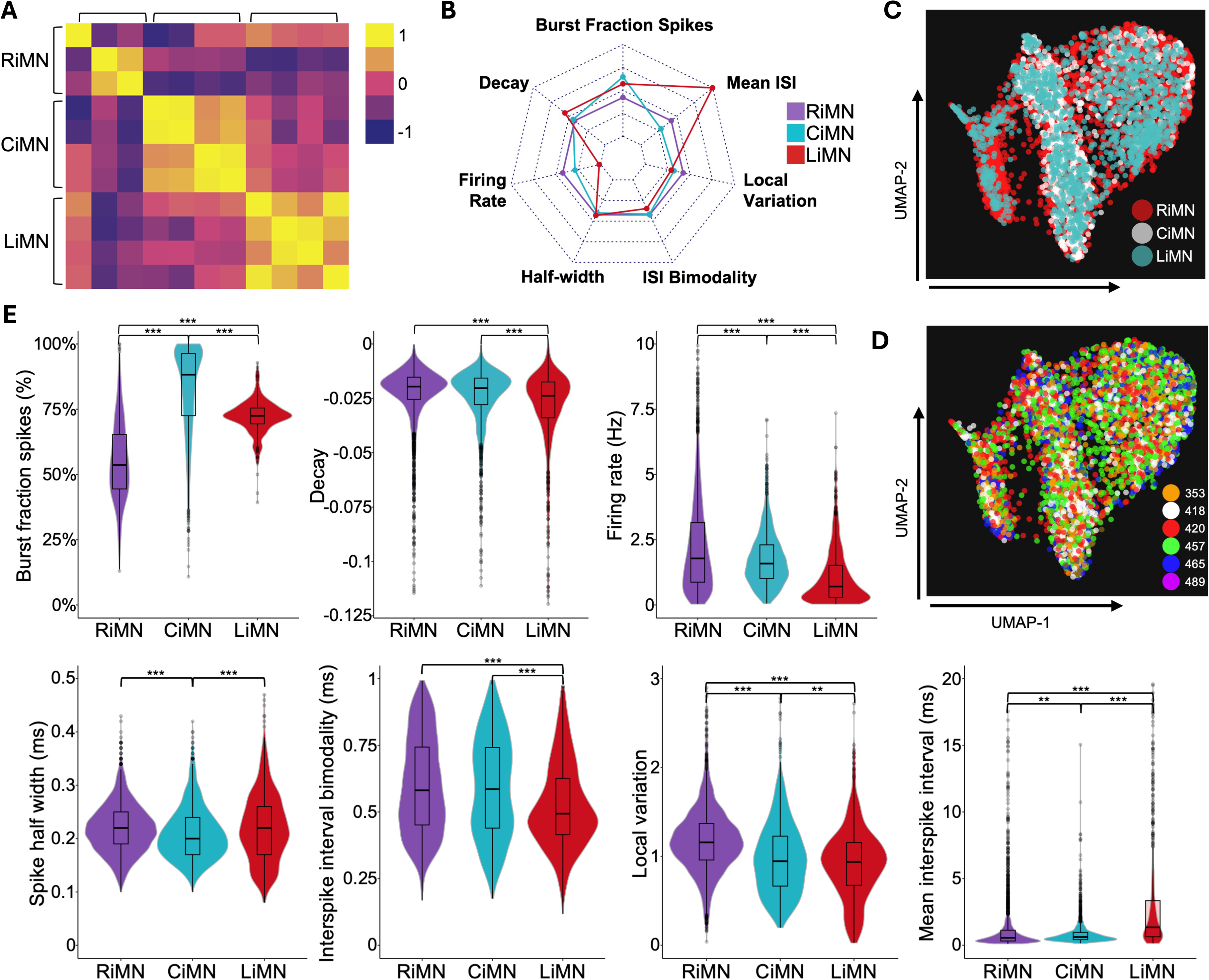
Regional iMN cultures exhibit distinct single-neuron electrophysiological profiles. Spike-sorted HD-MEA recordings were analyzed at day 45. **(A)** Pearson correlation matrix of well-level electrophysiological feature profiles across regional iMN populations, calculated using pairwise-complete observations. **(B)** Radar plot showing mean single-neuron electrophysiological feature values normalized to the corresponding RiMN means and expressed as fold changes, with RiMN set to 1 for each feature. **(C–D)** UMAP visualization of single-neuron feature profiles, colored by regional identity (C) or donor cell line (D). Each point represents one spike-sorted unit. **(E)** Violin plots with overlaid boxplots showing pooled unit-level distributions of burst fraction of spikes, waveform decay, firing rate, spike half-width, interspike interval bimodality, local variation, and mean interspike interval. Statistical comparisons were performed using one-way ANOVA followed by Tukey’s honestly significant difference test. Significance is indicated as *, P < 0.05; **, P < 0.01; and ***, P < 0.001. Data were obtained from three RiMN, four CiMN, and four LiMN cultures derived from independent iPSC lines. Only wells containing more than 100 units after quality control were included. diMN cultures exhibited insufficient activity for comparable single-neuron analysis. Missing feature values were excluded from the corresponding analyses; unit-level data are provided in Table S13.

UMAP visualization of single-neuron feature profiles showed overlapping distributions among the three regional populations, with CiMN units occupying a more restricted region of the embedding (Figure 6C). Units from different cell lines were interspersed throughout the embedding, suggesting that the observed electrophysiological structure was not explained by donor identity (Figure 6D). Pooled unit-level comparisons revealed differences across multiple electrophysiological features. Burst fraction of spikes was highest in CiMNs, intermediate in LiMNs, and lowest in RiMNs. Firing rates were highest in RiMNs and lowest in LiMNs, consistent with the overall pattern observed in the electrode-level measurements (Figures 6E and S7C). Mean ISIs were longest in LiMNs, and local variation was higher in RiMNs than in CiMNs and LiMNs. Burst fraction, firing rate, mean ISI, and local variation differed in all pairwise comparisons. CiMNs exhibited shorter spike half-widths than both other populations, whereas LiMNs showed lower ISI bimodality and more negative waveform decay values than RiMNs and CiMNs (Figure 6E). Notably, the higher burst fraction in CiMNs contrasted with the higher network burst peak firing rates observed in RiMNs (Figure S7C), highlighting differences between the fraction of spikes occurring within bursts and the intensity of peak network firing. Together, these analyses identify distinct electrophysiological profiles among regional iMN cultures.

## Discussion

In this study, we established three complementary protocols for generating human iPSC-derived MNs with controlled hindbrain, cervical, and lumbar identities. Across iPSC lines from six non-diseased donors balanced by sex, these protocols reproducibly generated regional HOX profiles while preserving MN identity and promoting region-appropriate subtype features. The resulting populations were transcriptionally distinct, showed the expected rostrocaudal relationships to marker-selected putative human fetal MNs, developed synchronized activity and electrically active axonal networks, and exhibited protocol-associated differences in single-neuron electrophysiological profiles. Regional patterning was achieved through a transient spheroid phase followed by adherent monolayer culture, providing regionally patterned iMNs in a format compatible with longitudinal imaging and HD-MEA analysis.

Conventional diMN differentiation reproduced progressive neural induction, collinear HOX activation, and MN specification, but generated broad and variable rostrocaudal programs. Similar heterogeneity was evident across the Answer ALS dataset, in which most iMN samples showed hindbrain/rhombomere or cervical features but varied substantially in posterior HOX expression. By controlling morphogen combinations and exposure windows, we shifted iMNs toward distinct positions along the rostrocaudal axis. LiMNs showed the greatest transcriptional divergence, consistent with their caudal identity and differentiation through a neuromesodermal progenitor-like intermediate. RiMNs and CiMNs remained more similar to diMNs because they shared more of the conventional differentiation framework, yet each acquired a distinct regional state. The resulting molecular profiles supported the intended identities: RiMNs suppressed cervical and posterior HOX programs and increased PHOX2B and ISL2 expression, supporting a hindbrain/rhombomere-associated state. CiMNs retained cervical HOX expression while restricting posterior programs and displayed a FOXP1-high, ALDH1A2-high, LHX1-maintained, and LHX3-low profile characteristic of cervical lateral motor column-like MNs. LiMNs activated posterior HOX genes across the HOXA, HOXC, and HOXD clusters while suppressing more rostral programs. Their preserved FOXP1 expression further indicated that caudalization remained compatible with limb-innervating MN-associated features. These populations should nevertheless be considered region-associated molecular states rather than fully resolved motor nuclei or columns, which will require single-cell and target-innervation analyses.

Comparison with human fetal spatial transcriptomic data provided an independent developmental reference. For direct benchmarking, putative fetal MNs were selected by detectable MNX1/HB9, ISL1, or ISL2 expression and aggregated into cervical, thoracic, and lumbar pseudobulk profiles. All-gene correlations were high within and across datasets, consistent with shared broad neural and MN transcriptional programs, but did not clearly distinguish rostrocaudal identity. HOX-restricted analysis was more informative and recovered the expected regional relationships: diMNs and CiMNs were most similar to the cervical reference, RiMNs showed progressively lower similarity to posterior regions, and LiMNs showed the greatest lumbar similarity among the iMN protocols. The main regional relationships were also preserved when a broader fetal reference selected using progenitor and neuronal/MN markers was compared using a combined marker and HOX gene panel. This agreement supports the regional interpretation across reference-selection strategies, while leaving the separate contributions of population selection and gene-panel expansion unresolved. The incomplete correspondence likely reflects overlapping fetal HOX boundaries, developmental and maturation differences, residual heterogeneity within the marker-selected fetal population, and technical differences between bulk and spatial transcriptomic measurements. These comparisons therefore support the direction and relative fidelity of regional patterning without implying complete equivalence to in vivo fetal MNs.

All three patterned populations developed synchronized firing and coordinated network bursts, whereas conventional diMNs showed comparatively sparse activity under the same HD-MEA conditions. RiMNs and CiMNs exhibited broad increases in network and axonal measurements, while LiMNs developed synchronized activity and enhanced axonal propagation-related properties. Axon-tracking further revealed extended, branched, and electrically active neuronal processes, demonstrating the formation of connected networks capable of signal propagation and coordination. These functional observations were complemented by NEFH-based morphological quantification, which demonstrated greater skeleton length and more network junctions per nucleus in RiMNs and CiMNs, thus providing complementary structural evidence for their enhanced functional network activity.

Spike-sorted analyses provided complementary functional characterization, revealing greater similarity in combined single-neuron feature profiles within regional protocols than across protocols (Figure 6A). LiMN units exhibited lower firing rates and longer mean interspike intervals, whereas CiMN units showed a higher fraction of spikes occurring within bursts (Figures 6B and 6E). These differences complement the regional HOX programs, supporting molecular and functional distinctions among the cultures. However, overlapping unit-level profiles and variation across cell lines indicate heterogeneous electrophysiological states within regional cultures rather than uniformly separable neuronal populations (Figures 6C, 6D, and S8). Regional differences therefore emerge from the combined feature profiles alongside substantial variation among individual units. The greater firing activity of RiMNs and CiMNs relative to LiMNs and diMNs may have several non-mutually exclusive explanations. Regional MN populations may differ intrinsically in excitability, synaptic connectivity, or network-assembly programs^48,49^, although the present data cannot separate these properties from protocol-dependent effects. The four populations also follow distinct developmental trajectories, such that cultures analyzed on the same differentiation day may not represent equivalent stages of functional maturation^18,49^, particularly for LiMNs generated through a neuromesodermal progenitor-like intermediate. Differences in cellular composition, including the abundance of glial cells, residual progenitors, or excitatory and inhibitory neuronal populations, could further influence network firing and synchronization^46,50^. Distinguishing among these possibilities will require integrated molecular and electrophysiological analyses that link recorded units to defined cell types and maturation states. The transient spheroid phase may contribute to these properties by promoting cell-cell interactions, early network organization, and activity-dependent maturation, although the present study does not separate these effects from those of morphogen patterning. Importantly, the protocols ultimately yield adherent monolayers rather than terminal spheroid or organoid cultures. This endpoint facilitates morphological tracking, electrophysiological recording, single-cell manipulation, and axon tracking and supports broader implementation across laboratories and downstream applications.

The reproducible regional profiles generated here provide an opportunity to investigate the biochemical functions of individual HOX proteins during human MN development. Comparing the same perturbation across RiMN, CiMN, and LiMN backgrounds could reveal how regional context influences HOX activity, cofactor interactions, cross-repressive HOX networks, MN subtype specification, axonal growth, and excitability. A key next step will be to apply time-resolved single-cell RNA sequencing across differentiation to define the cellular composition of each culture, resolve intermediate states and developmental trajectories, and identify region-specific transcriptional programs and downstream HOX targets. These protocols also provide region-matched populations for investigating selective MN degeneration. Generating RiMNs, CiMNs, and LiMNs from the same genetic background would allow regional susceptibility to be compared without confounding differences between donors or differentiation platforms. Application to disease-associated and gene-edited isogenic lines could then distinguish shared mechanisms from changes affecting specific MN subtypes or anatomical levels, while single-cell resolution could help separate altered cell-state composition from cell-intrinsic disease signatures. Together, these approaches could support longitudinal disease modeling, mechanistic studies, and region-informed therapeutic screening.

Several limitations remain. Bulk RNA-seq cannot resolve residual cellular heterogeneity or determine whether regional markers coexist within individual cells. Although the fetal reference was enriched for putative MNs through MNX1/HB9, ISL1, or ISL2 expression, this marker-based selection does not provide a purified or subtype-resolved MN population and captures only one developmental stage. HD-MEA activity also does not establish mature motor output or neuromuscular connectivity. Co-culture with glial cells, region-appropriate muscle targets, neuromuscular junction analysis, and in vivo integration studies will be needed to assess target-specific connectivity and motor output. Overall, this three-protocol pipeline provides a reproducible framework for investigating how rostrocaudal developmental programs shape human MN identity, function, and selective vulnerability.

## Resource availability

### Lead contact

Further information and requests for resources and reagents should be directed to and will be fulfilled by the lead contact, **[Ritchie Ho]**.

### Materials availability

This study did not generate unique genetically modified cell lines or proprietary reagents. The regional differentiation protocols and associated analytical workflows are described below.

### Data and code availability

Bulk RNA-seq data generated in this study have been deposited in the NCBI Gene Expression Omnibus (GEO) under accession number GSE349145. The processed expression matrices and metadata required to reproduce the reported analyses will be made available with the deposited dataset. Code for RNA-seq analysis and figure generation is available at https://github.com/ningken14/Regional-iMN-RNAseq, and code for HD-MEA data processing and analysis is available at https://github.com/ianTindel/SpikeInterfaceAndDeephysPipeline. Publicly available Answer ALS transcriptomic data and the Lawrence et al. (2024) week 9 fetal spatial transcriptomic dataset were accessed as described below. Any additional information required to reanalyze the data is available from the lead contact upon reasonable request.

## Experimental model and subject details

### Human iPSC lines

Experiments used six age-matched human iPSC lines derived from non-diseased donors, comprising three female-derived and three male-derived lines. Individual lines were treated as independent biological replicates; line-specific metadata are summarized in Table S1.

### Sex as a biological variable

This study included six iPSC lines derived from non-diseased donors, comprising three female-derived and three male-derived lines matched for donor age. Lines from both sexes were represented across the regional differentiation protocols and downstream molecular and functional analyses. This balanced design was used to evaluate protocol reproducibility across donor backgrounds and minimize sex-related sampling bias. However, the study was not designed or powered to identify sex-dependent differences, and no conclusions regarding effects of donor sex were drawn.

### iPSC quality control

iPSC cultures were routinely assessed for characteristic colony morphology and the absence of spontaneous differentiation. All lines were confirmed to be free of mycoplasma and to meet institutional criteria for pluripotency and genomic integrity before experimental use. Only cultures at or below passage 45 were used in this study.

## Method details

### iPSC maintenance

iPSCs were maintained on tissue culture-treated plates coated with growth factor-reduced Matrigel (Corning, catalog no. 354230) in mTeSR Plus medium (STEMCELL Technologies, catalog no. 100-0276) at 37°C in a humidified atmosphere containing 5% CO₂. Cultures were passaged every 5–7 days at approximately 75%–85% confluency using Versene solution (Gibco, catalog no. 15040066) and routinely reseeded at a 1:10 split ratio. Only cultures meeting the quality-control criteria described above were used for differentiation.

### Conventional diMN differentiation

Conventional diMNs were generated using an established three-stage adherent differentiation protocol^9,19^. iPSCs were dissociated with Accutase (STEMCELL Technologies, catalog no. 07920) and seeded at 2 × 10⁵ cells per well on Matrigel-coated 6-well plates in mTeSR Plus supplemented with 10 μM Y-27632 (Selleckchem, catalog no. S1049). After 24 h, differentiation was initiated (day 0) by replacing mTeSR Plus with Stage 1 medium composed of 47.5% IMDM (Gibco, catalog no. 12440053), 47.5% F12 (Gibco, catalog no. 11765062), 1% non-essential amino acids (Gibco, catalog no. 11140050), 2% B27 supplement with vitamin A (Gibco, catalog no. 17504044), 1% N2 supplement (Gibco, catalog no. 17502048), and 1% antibiotic-antimycotic (Gibco, catalog no. 15240062). Stage 1 medium was supplemented with 0.2 μM LDN193189 (Bio-Techne, catalog no. 6053), 3 μM CHIR99021 (Cayman Chemical, catalog no. 13122), and 10 μM SB431542 (Cayman Chemical, catalog no. 13031). Cultures received fresh Stage 1 medium daily from days 0–5 to induce neuroepithelial identity. On day 6, neuroepithelial cells were dissociated with Accutase and replated at 7.5 × 10⁵ cells per well on Matrigel-coated 6-well plates in Stage 2 platedown medium. Stage 2 medium contained the same basal components and concentrations of LDN193189, CHIR99021, and SB431542 as Stage 1 medium, together with 0.1 μM all-trans retinoic acid (Stemgent, catalog no. 04-0021) and 1 μM SAG (Cayman Chemical, catalog no. 11914) to promote rostrocaudal and ventral patterning. The platedown medium was additionally supplemented with 10 μM Y-27632. On day 7, cultures were transferred to Stage 2 medium without Y-27632, followed by approximately 75% medium replacement on days 9 and 11. Terminal MN maturation was initiated on day 12 using Stage 3 medium composed of 47.5% IMDM, 47.5% F12, 1% non-essential amino acids, 2% B27 supplement with vitamin A, 1% N2 supplement, and 1% antibiotic-antimycotic. Stage 3 medium was supplemented with 0.1 μM SAG, 0.5 μM all-trans retinoic acid, 0.1 μM dibutyryl-cAMP (STEMCELL Technologies, catalog no. 73884), 0.1 μM Compound E (Millipore, catalog no. 565790), 2.5 μM DAPT (Sigma-Aldrich, catalog no. D5942), 200 ng/mL ascorbic acid (Sigma-Aldrich, catalog no. A4403), 10 ng/mL BDNF (PeproTech, catalog no. 450-02), and 10 ng/mL GDNF (PeproTech, catalog no. 450-10). Approximately 75% of the medium was replaced every other day through day 30, and mature diMNs were collected or analyzed from day 32 onward. Cultures were maintained in an adherent monolayer format throughout differentiation, with replating at the transition from neuroepithelial cells to MN progenitors.

### Transient 3D-to-2D diMN control (diMN 232D)

For the diMN 232D control, cells received the same basal media and morphogen sequence as conventional diMNs but were transiently cultured as spheroids during the progenitor stage. On day 6, cells were dissociated with Accutase and seeded at 5 × 10⁵ cells per Cell Spheroid Honeycomb Culture Chip (Suzhou Vivoid Biotechnology, catalog no. AUT042801). Spheroids were maintained until day 12, dissociated with Accutase, and replated as adherent cultures. Cells were subsequently matured under the same conditions used for conventional diMNs.

### RiMN differentiation

RiMNs were generated using the optimized RiMN-E differentiation protocol. Basal medium (BM2) comprised 47% DMEM/F12 without L-glutamine, 47% Neurobasal medium, 1% non-essential amino acids, 2% B27 with vitamin A, 1% N2, 1% antibiotic-antimycotic, and 1% GlutaMAX. On day −1, iPSCs were dissociated with Accutase and seeded at 2 × 10⁵ cells per well in 6-well tissue-culture-treated plates containing mTeSR Plus supplemented with 10 μM Y-27632. Differentiation was initiated on day 0 by replacing the maintenance medium with RiMN-E Stage 1 medium, consisting of BM2 supplemented with 1.5 μM CHIR99021, 0.5 μM LDN193189, and 10 μM SB431542. Cultures were maintained under neural induction conditions through day 5. On day 6, cells were dissociated with Accutase and seeded at 1 × 10⁵ cells per Cell Spheroid Honeycomb Culture Chip (Suzhou Vivoid Biotechnology, catalog no. AUT042801) in RiMN-E Stage 2 medium supplemented with 10 μM Y-27632 to generate progenitor spheroids. From days 6–11, spheroids were maintained in Stage 2 medium containing BM2 supplemented with 1.5 μM CHIR99021, 0.5 μM LDN193189, 10 μM SB431542, 1 μM SAG, 20 ng/mL EGF (Thermo Fisher Scientific, catalog no. AF-100-15), 10 ng/mL FGF8b (Thermo Fisher Scientific, catalog no. 100-25), and 0.1 μM all-trans retinoic acid to promote ventral hindbrain/rhombomere-associated patterning. On day 12, patterned spheroids were dissociated with Accutase and replated as adherent cultures at 5 × 10⁶ cells per well on Matrigel-coated tissue-culture-treated plates. From days 12–32, cultures were maintained in RiMN-E Stage 3 medium, comprising BM2 supplemented with 2.5 μM DAPT, 0.1 μM dibutyryl-cAMP, 0.1 μM Compound E, 200 ng/mL ascorbic acid, 10 ng/mL BDNF, 10 ng/mL GDNF, 0.1 μM SAG, and 0.1 μM all-trans retinoic acid. Medium was replaced every other day throughout maturation, and mature RiMNs were collected or analyzed from day 32 onward.

### CiMN differentiation

CiMNs were generated using the optimized CiMN-H differentiation protocol. Basal medium (BM2) comprised 47% DMEM/F12 without L-glutamine, 47% Neurobasal medium, 1% non-essential amino acids, 2% B27 with vitamin A, 1% N2, 1% antibiotic-antimycotic, and 1% GlutaMAX. On day −1, iPSCs were dissociated with Accutase and seeded at 2 × 10⁵ cells per well in 6-well tissue-culture-treated plates containing mTeSR Plus supplemented with 10 μM Y-27632. Differentiation was initiated on day 0 by replacing the maintenance medium with CiMN-H Stage 1 medium, consisting of BM2 supplemented with 3.0 μM CHIR99021, 0.5 μM LDN193189, and 10 μM SB431542. Cultures were maintained under neural induction conditions through day 5. On day 6, cells were dissociated with Accutase and seeded at 1 × 10⁵ cells per Cell Spheroid Honeycomb Culture Chip (Suzhou Vivoid Biotechnology, catalog no. AUT042801) in CiMN-H Stage 2 medium supplemented with 10 μM Y-27632 to generate progenitor spheroids. From days 6–11, spheroids were maintained in Stage 2 medium containing BM2 supplemented with 3.0 μM CHIR99021, 0.5 μM LDN193189, 10 μM SB431542, 1 μM SAG, 20 ng/mL EGF (Thermo Fisher Scientific, catalog no. AF-100-15), and 0.1 μM all-trans retinoic acid to promote ventral cervical/brachial-associated patterning. On day 12, patterned spheroids were dissociated with Accutase and replated as adherent cultures at 5 × 10⁶ cells per well on Matrigel-coated tissue-culture-treated plates. From days 12–32, cultures were maintained in CiMN-H Stage 3 medium, comprising BM2 supplemented with 2.5 μM DAPT, 0.1 μM dibutyryl-cAMP, 0.1 μM Compound E, 200 ng/mL ascorbic acid, 10 ng/mL BDNF, 10 ng/mL GDNF, 0.1 μM SAG, and 1.0 μM all-trans retinoic acid. Medium was replaced every other day throughout maturation, and mature CiMNs were collected or analyzed from day 32 onward.

### LiMN differentiation

LiMNs were generated using the optimized protocol through a neuromesodermal progenitor-like intermediate followed by caudal and ventral patterning. Basal medium 1 (BM1) comprised 47% DMEM/F12 without L-glutamine, 47% Neurobasal medium, 1% non-essential amino acids, 2% B27 without vitamin A (Gibco, catalog no. 12587010), 1% N2, 1% antibiotic-antimycotic, and 1% GlutaMAX. Basal medium 2 (BM2) contained the same components but used B27 with vitamin A. On day -5, iPSCs were dissociated and seeded at 5 × 10^5^ cells per Cell Spheroid Honeycomb Culture Chip (Suzhou Vivoid Biotechnology, catalog no. AUT042801) in NMP-1 medium, comprising BM1 supplemented with 20 ng/mL FGF2 (Thermo Fisher Scientific, catalog no. 100-18B) and 10 μM Y-27632. Cultures were maintained in NMP-1 medium through day -3 and then in NMP-2 medium, comprising BM1 supplemented with 20 ng/mL FGF2 and 3 μM CHIR99021, from days -2 to -1 to induce a neuromesodermal progenitor-like state. On day 0, spheroids were removed from the chips and transferred to ultra-low-attachment plates (Corning, catalog no. 3473). From days 0–3, spheroids were maintained in LiMN-M3 medium, comprising BM1 supplemented with 3 μM CHIR99021, 0.2 μM LDN193189, and 10 μM SB431542. On day 4, cultures were changed to LiMN-M4 medium containing BM1 supplemented with 3 μM CHIR99021, 0.2 μM LDN193189, 150 ng/mL FGF8b, and 50 ng/mL GDF11 (Thermo Fisher Scientific, catalog no. 120-11). On day 5, CHIR99021 and LDN193189 were withdrawn, and spheroids were maintained in LiMN-M5 medium containing BM1 supplemented with 150 ng/mL FGF8b and 50 ng/mL GDF11. To control spheroid size, aggregates were mechanically chopped on day 6 and returned to suspension culture. From days 6–11, spheroids were cultured in LiMN-M9 medium, comprising BM2 supplemented with 0.1 μM all-trans retinoic acid, 1 μM SAG, and 50 ng/mL GDF11, to promote caudal and ventral MN patterning. On day 12, spheroids were dissociated with Accutase and replated at 5 × 10^6^ cells per well on Matrigel-coated tissue-culture-treated plates. From days 12–21, cultures were maintained in LiMN-M11 medium, comprising BM1 supplemented with 2.5 μM DAPT, 0.1 μM dibutyryl-cAMP, 0.1 μM Compound E, 200 ng/mL ascorbic acid, 10 ng/mL BDNF, 10 ng/mL GDNF, and 25 ng/mL GDF11. From days 22–32, cultures were maintained in LiMN-M12 medium, comprising BM2 supplemented with 2.5 μM DAPT, 0.1 μM dibutyryl-cAMP, 0.1 μM Compound E, 200 ng/mL ascorbic acid, 10 ng/mL BDNF, 10 ng/mL GDNF, and 0.1 μM all-trans retinoic acid. Medium was replaced every other day throughout adherent maturation, and mature LiMNs were collected or analyzed from day 32 onward.

### Spheroid imaging and morphological analysis

RiMN, CiMN, and LiMN spheroids were imaged longitudinally in brightfield using an EVOS M5000 Imaging System. Images were acquired at the differentiation time points indicated in Figure S1 and analyzed using ImageJ/Fiji. Following spatial calibration and segmentation of the spheroid boundary, projected area was calculated from the enclosed region. Roundness was determined using the Fiji Shape Descriptors function, with values approaching 1 indicating greater circularity. Feret’s diameter, defined as the maximum caliper distance between any two points along the segmented boundary, was used as an additional measure of spheroid size. Measurements were tracked longitudinally for each independent iPSC line, with each colored trajectory in Figure S1 representing a separate line.

### RNA isolation, reverse transcription, and quantitative PCR

Total RNA was isolated using the RNeasy Mini Kit (Qiagen, catalog no. 74104) with on-column digestion using the RNase-Free DNase Set (Qiagen, catalog no. 79256). RNA concentration and purity were assessed using a NanoDrop One spectrophotometer. First-strand cDNA was synthesized using the ProtoScript II First Strand cDNA Synthesis Kit (New England Biolabs, catalog no. E6560L) according to the manufacturer’s instructions. Quantitative PCR was performed using SYBR Green chemistry (Applied Biosystems, catalog no. 4309155) on a Bio-Rad CFX384 Real-Time PCR Detection System. Reactions were run in technical triplicate for 40 amplification cycles. Gene expression was normalized to GAPDH and calculated using the 2^−ΔCt method in GraphPad Prism or R. Primer sequences are listed in Table S2.

### Immunofluorescence staining and image quantification

Cultures were fixed with 4% paraformaldehyde for 15 min at room temperature and washed three times with 1× PBS. Cells were permeabilized in 0.3% PBST and blocked in 5% normal donkey serum in 0.1% PBST. Primary antibodies were diluted in blocking solution and incubated overnight at 4°C. After three washes with 1× PBS, species-appropriate fluorophore-conjugated secondary antibodies were applied for 1 h at room temperature protected from light. Nuclei were counterstained with DAPI. Antibody details are listed in Table S3.

Images were acquired using a Nikon AX R confocal microscope with NSPARC detection under constant acquisition settings across conditions within each experiment. For HOX and other nuclear markers, DAPI-positive nuclei were segmented in Fiji, marker-positive fractions were calculated relative to total DAPI-positive nuclei, and background-subtracted mean fluorescence intensity was measured. HOXC6 and HOXC10 marker-positive fractions were analyzed using ordinary one-way ANOVA followed by Tukey’s multiple-comparisons test. HOXC10 mean fluorescence intensity was compared between diMNs and LiMNs using a t-test. ISL1-positive fractions and mean fluorescence intensities were compared between each regional iMN population and diMNs using Welch’s t-test. For NEFH morphological analysis, maximum-intensity projections of confocal Z-stacks were segmented and skeletonized using an automated workflow to quantify skeleton length, 2D network junctions, local orientation dispersion, and radial alignment; skeleton length and junction counts were normalized to DAPI-positive nuclei. Three images per cell line and condition were averaged to obtain one cell-line-level value. NEFH data were analyzed using ordinary one-way ANOVA followed by Dunnett’s multiple-comparisons test, comparing each regional iMN population with diMNs. Data visualization and statistical analyses were performed using GraphPad Prism.

### Bulk RNA sequencing and preprocessing

RNA was collected from iPSC, diMN, RiMN, CiMN, and LiMN cultures generated across two experimental batches. Total RNA was isolated using the RNeasy Mini Kit with on-column RNase-Free DNase treatment (Qiagen, catalog nos. 74104 and 79256), and RNA concentration and purity were assessed using a NanoDrop One spectrophotometer. Library preparation and sequencing were performed by Plasmidsaurus. FASTQ quality was assessed using FastQC v0.12.1. Reads were processed using fastp v0.24.0 with poly-X and 3′ quality trimming, a minimum Phred score of 15, and a minimum retained length of 50 bp. Reads were aligned using STAR v2.7.11, coordinate-sorted with samtools v1.22.1, and deduplicated using UMIcollapse v1.1.0. Alignment and library quality were evaluated using RSeQC v5.0.4 and Qualimap v2.3 and summarized with MultiQC v1.32. Strand-specific gene counts were generated using featureCounts v2.1.1, with fractional assignment of multimapping reads and aggregation by gene_id. Count matrices from both batches were harmonized by gene identifier and symbol, merged by gene_id, and deduplicated. The original batch-1 457_RiMN sample was excluded as a quality-control outlier and replaced by a batch-2 sample. diMNv1 and preliminary batch-1 CiMN-E samples were excluded, and the optimized CiMN-H condition was analyzed as CiMN. Batch-2 diMN, RiMN, and LiMN samples from cell line 418 were retained as bridge samples for estimation of batch effects. Counts were TMM-normalized, and genes with >1 count per million in at least three samples were retained for limma-voom transformation. Batch- and cell-line-adjusted expression values were used for visualization and PCA, whereas differential expression models incorporated protocol, batch, and cell line as covariates.

### Normalization, principal component analysis, and loading-gene enrichment

Counts were analyzed in R using edgeR and limma. Genes with counts per million (CPM) greater than 1 in at least three samples were retained. Trimmed mean of M values (TMM) normalization factors were calculated with edgeR, and voom transformation was performed with a design containing sequencing batch and protocol. For visualization, batch and cell-line effects were removed using limma::removeBatchEffect while preserving protocol-associated variation. Principal component analysis (PCA) was performed with prcomp on the transposed corrected voom logCPM matrix without additional feature scaling. Ninety-five percent normal ellipses were displayed for groups with sufficient replicates. A second PCA was refitted after restricting the data to diMN, RiMN, and CiMN samples to resolve differences among protocols sharing a similar differentiation backbone. The top positive and negative loading genes for PC1 and PC2 were visualized by heatmap. Gene Ontology (GO) Biological Process over-representation analysis was performed with clusterProfiler after mapping gene symbols to Entrez identifiers using org.Hs.eg.db. The expressed-gene set served as the background universe, and Benjamini–Hochberg correction was applied. The full PCA loading analysis used the top 250 genes from each loading direction, with displayed GO terms ranked by fold enrichment. The restricted PC2 analysis also used the top 250 genes from each direction; the eight terms with the lowest adjusted P-values were displayed using signed −log₁₀(adjusted P-value).

### Differential expression and pathway enrichment analysis

Genes were filtered using edgeR::filterByExpr with protocol as the grouping factor and normalized by the trimmed mean of M values method. Voom transformation and linear modeling were performed using a no-intercept design incorporating protocol, sequencing batch, and cell line. Pairwise protocol contrasts were fitted with limma and subjected to empirical Bayes moderation. Differentially expressed gene sets used for volcano-plot classification and overlap analyses were defined by a Benjamini–Hochberg-adjusted P-value < 0.05 and an absolute log₂ fold change > 1. Tables S10–S12 provide the complete lists of genes with a Benjamini–Hochberg-adjusted P-value < 0.05 for RiMN, CiMN, and LiMN comparisons with diMN, respectively. Rows retained only as Ensembl-style identifiers without an interpretable gene symbol were omitted from displayed volcano plots and enrichment analyses. For Gene Ontology (GO) Biological Process over-representation analysis, upregulated and downregulated gene sets were selected separately using a Benjamini–Hochberg-adjusted P-value < 0.1 and an absolute log₂ fold change > 1. Enrichment analysis was performed using clusterProfiler, with gene symbols mapped to Entrez identifiers using org.Hs.eg.db. Mapped expressed genes retained in the fitted model were used as the background universe. GO terms were retained using P-value, Benjamini–Hochberg-adjusted P-value, and q-value cutoffs of 0.05. Fold enrichment was calculated as the ratio of GeneRatio to BgRatio. Up to six terms with the highest fold enrichment were displayed for each direction, without exclusions based on term descriptions. For visualization, fold enrichment was assigned a positive sign for upregulated gene sets and a negative sign for downregulated gene sets. DEG overlaps were visualized using UpSetR, with gene sets defined by a Benjamini– Hochberg-adjusted P-value < 0.05 and an absolute log₂ fold change > 1.

### Expression heatmaps and targeted gene comparisons

Heatmaps were generated from batch- and cell-line-corrected voom logCPM values. Where indicated, expression was averaged by protocol and standardized gene-wise as z-scores. HOXA, HOXB, HOXC, and HOXD genes were ordered by paralog number. Targeted gene comparisons included only cell lines represented in both conditions, with replicate observations averaged within each cell line and protocol. Two-sided paired t-tests were performed on within-cell-line differences, and P-values were adjusted within each predefined marker-analysis set using the Benjamini–Hochberg procedure. Significance brackets display categories derived from the adjusted P-values.

### Answer ALS transcriptomic analysis

To assess rostrocaudal heterogeneity among conventionally generated iMNs, a transcriptomic data matrix of 508 iPSC-derived MN samples was obtained from the Answer ALS Data Portal (https://dataportal.answerals.org/home). Ensembl identifiers were mapped to HGNC gene symbols using AnnotationDbi and org.Hs.eg.db; genes without mapped symbols were removed, and the first entry was retained when multiple Ensembl identifiers mapped to the same symbol. Raw counts were processed using DESeq2 with an intercept-only design, and expression values were transformed using the blind variance-stabilizing transformation without additional batch correction. HOX genes with non-zero variance were selected and visualized as sample-level distributions. Genes were arranged a priori from rostral hindbrain/rhombomere-associated genes through cervical/brachial and thoracic genes to caudal/lumbar genes rather than by unsupervised clustering. Replicate libraries were retained as separate sample-level observations.

### Human fetal spatial transcriptomic analysis

Processed week 9 cervical, thoracic, and lumbar spatial transcriptomic datasets were downloaded as H5AD files from the CellGenI spinal development data portal (https://spinal-development.cellgeni.sanger.ac.uk/) accompanying Lawrence et al. (2024). Files were imported into R using zellkonverter::readH5AD, converted to Seurat objects, and merged. Count matrices stored within the objects were used for downstream analysis, and gene symbols were harmonized and made unique. Data were log-normalized using a scale factor of 10,000. The 3,000 most variable features were selected using the vst method, followed by scaling, PCA, and UMAP based on the first 30 principal components. Numeric spatial clusters were annotated using markers reported in the original study and markers identified within the week 9 dataset. Positive cluster markers were calculated using Seurat::FindAllMarkers with a minimum detection fraction of 0.10 and a log₂ fold-change threshold of 0.25. The top 30 markers from each cluster were compared with canonical lineage markers and the marker reference provided by the original study. Curated annotations included chondrocyte/chondroprogenitor, muscle/myocyte, glial/Schwann-like, cord neuron/GABAergic neuron, sensory neuron/nociceptor, sensory neuron/early nociceptor, erythrocyte, and tenocyte/fibrous mesenchyme. Putative fetal MN profiles were selected by detectable expression of MNX1, ISL1, or ISL2; the expanded comparison in Figure S5F used a broader panel of progenitor and neuronal/MN markers. Counts were aggregated by anatomical region, TMM-normalized using edgeR, and converted to logCPM with a prior count of 2. Regional iMN expression was averaged by protocol using batch-and cell-line-corrected voom logCPM values. Spearman correlations were calculated across shared genes using all-gene, HOX-only, or combined marker–HOX panels, with pairwise complete observations. CHAT was excluded from marker-based correlations because it was not detectably expressed in the fetal reference. Correlation matrices were displayed without hierarchical reordering.

### High-density multielectrode array recordings

Neuronal network activity and axonal signal propagation were measured using a MaxWell Biosystems high-density multielectrode array (HD-MEA) platform according to the manufacturer’s protocol. On day 12, cultures were dissociated and plated at 10,000–15,000 cells/mm² onto MaxWell six-well HD-MEA plates coated with 0.07% polyethylenimine (Sigma-Aldrich, catalog no. P3143) and 5 μg/mL laminin-521 (BioLamina, catalog no. LN521). Cells were maintained in BrainPhys Neuronal Medium (STEMCELL Technologies, catalog no. 05790), with partial medium replacement every 2 days before and throughout longitudinal recording. Recordings were acquired using the MaxTwo system in a temperature- and CO₂-controlled environment. Longitudinal measurements were performed at the differentiation days indicated in Figures 5 and S6–S7. Cross-protocol comparisons at approximately day 45 used one summary value per independent iPSC line.

### Spike sorting and single-neuron analysis

HD-MEA recordings acquired at day 45 were decompressed from .h5 files using the MaxWell Biosystems decompression library in MATLAB according to the manufacturer’s documentation. Spike sorting was performed using the Tridesclous2 sorter implemented in the SpikeInterface Python package, and outputs were saved in NumPy (.npy) format^51,52^. Spike-sorted recordings were processed separately for each well using the DeePhys MATLAB package^52,53^, including quality control to exclude low-quality signals, identification of putative single-neuron units, and extraction of electrophysiological features. Wells containing more than 100 units after quality control were retained, and their feature datasets were aggregated at the plate level. Analyzable units were obtained from three RiMN, four CiMN, and four LiMN cultures, whereas conventional diMN cultures exhibited insufficient activity for comparable single-neuron analysis. Seven unit-level features were examined: burst fraction of spikes, waveform decay, firing rate, spike half-width, interspike interval (ISI) bimodality, local variation, and mean ISI. Pooled unit-level feature values were compared among regional iMN populations using one-way ANOVA followed by Tukey’s multiple-comparisons test. Similarity between well-level electrophysiological feature profiles was assessed using Pearson correlation with pairwise-complete observations. Individual feature distributions were compared between wells using Kolmogorov–Smirnov tests, with P-values adjusted for multiple comparisons using the Benjamini–Hochberg procedure.

### Quantification and statistical analysis

Independent iPSC lines were treated as biological replicates. Multiple images, wells, spheroids, or HD-MEA recordings obtained from the same cell line and condition were not considered independent biological replicates and were summarized at the cell-line level unless otherwise specified. Exact sample sizes, differentiation days, statistical tests, and error-bar definitions are provided in the corresponding figure legends. Samples were excluded only according to the prespecified experimental or sequencing quality-control criteria described above. Bulk RNA-seq differential expression was assessed using limma-voom models incorporating protocol, sequencing batch, and cell line, with Benjamini–Hochberg correction for multiple testing. Targeted expression comparisons were performed using two-sided paired t-tests across matched cell lines, with Benjamini–Hochberg correction within the relevant gene–contrast family. qPCR, immunofluorescence, and spheroid measurements were analyzed using GraphPad Prism or R, with the tests applied to each experiment specified in the figure legends. Unless otherwise indicated, P or adjusted P < 0.05 was considered significant. Significance is denoted as ns, P ≥ 0.05; *, P < 0.05; **, P < 0.01; ***, P < 0.001; and ****, P < 0.0001.

### Software and algorithms

|  |  |  |
| --- | --- | --- |
| R | 4.6.0 | Statistical computing |
| edgeR | 4.10.1 | Gene filtering, TMM normalization, and pseudobulk processing |
| limma | 3.68.3 | Voom transformation, linear modeling, and batch/cell-line correction |
| DESeq2 | 1.52.0 | Count-based expression analysis and variance-stabilizing transformation |
| clusterProfiler | 4.20.0 | Gene Ontology enrichment analysis |
| org.Hs.eg.db | 3.18.0 | Human gene annotation |
| AnnotationDbi | 1.74.0 | Gene identifier mapping |
| biomaRt | 2.68.0 | Gene annotation utilities |
| Seurat | 5.5.0 | Spatial transcriptomic processing, UMAP, and marker analysis |
| SeuratObject | 5.4.0 | Seurat data structures and assay-layer handling |
| SingleCellExperiment | 1.34.0 | Single-cell and spatial expression data structures |
| SummarizedExperiment | 1.42.0 | Expression assays and associated metadata |
| zellkonverter | 1.22.0 | H5AD import and conversion |
| Matrix | 1.7-5 | Sparse matrix operations |
| pheatmap | 1.0.13 | Expression and loading heatmaps |
| UpSetR | 1.4.1 | DEG-overlap plots |
| ggplot2 | 4.0.3 | Data visualization |
| ggpubr | 0.6.3 | Statistical annotations and publication-oriented plots |
| ggrepel | 0.9.8 | Non-overlapping plot labels |
| patchwork | 1.3.2 | Multipanel figure assembly |
| ggsignif | 0.6.4 | Statistical significance annotations |
| RColorBrewer | 1.1-3 | Plot color palettes |
| scales | 1.4.0 | Plot scales and label formatting |
| dplyr | 1.2.1 | Data manipulation and summarization |
| tidyr | 1.3.2 | Data reshaping |
| tibble | 3.3.1 | Tabular data structures |
| readr | 2.2.0 | Delimited text-file import |
| readxl | 1.5.0 | Excel-file import |
| stringr | 1.6.0 | String processing |
| purrr | 1.2.2 | Functional programming and list operations |
| plyr | 1.8.9 | Data manipulation utilities |
| reshape2 | 1.4.5 | Data reshaping utilities |
| broom | 1.0.13 | Statistical result formatting |
| rstatix | 0.7.3 | Statistical testing utilities |
| tidyverse | 2.0.0 | Data manipulation and visualization package collection |
| MaxWell Biosystems / MaxLab Live | 25.1.8.1 | HD-MEA acquisition and axon tracking |
| GraphPad Prism | 10.3.0 (461) | Wet-laboratory statistics and plots, where applicable |
| ImageJ / Fiji | 1.54p | Spheroid and immunofluorescence quantification |
| MATLAB | R2026a | HD-MEA data decompression and electrophysiological analysis |
| DeePhys | 2.0.0 | Spike-sorted unit quality control and electrophysiological feature extraction |
| SpikeInterface | 0.104.8 | Spike sorting using Tridesclous2 |

## Disclosure

AI tools assisted with language refinement, code debugging and organization, and selected data organization tasks under author supervision. The authors reviewed the resulting text, code, and data and retain full responsibility for the scientific content, analyses, and conclusions.

## Author Contributions

RH and KN conceptualized and designed the study. KN, IT, BKS, BZ and RH performed experiments. IT analyzed HD-MEA data. KN carried out data processing and analysis. RH supervised all aspects of the study. RH, KN, and IT wrote the manuscript, with editing input from BKS.

## Acknowledgements

We thank the Cedars-Sinai Biomanufacturing Center for providing iPSC cell lines. RH was supported by grants from the National Institute on Aging (R00AG056678), Winnick Foundation Research Scholars Award, California Institute for Regenerative Medicine Discovery-2 (CIRM-DISC2 15010). KN and BS were supported by the California Institute for Regenerative Medicine (EDUC4-12751). Data used in the preparation of this article were obtained from the ANSWER ALS Data Portal (AALS-01184). For up-to-date information on the study, visit https://dataportal.answerals.org.

## Notes

### Competing Interest Statement

The authors have declared no competing interest.

## References

1 Dasen, J. S., Tice, B. C., Brenner-Morton, S. & Jessell, T. M. A Hox regulatory network establishes motor neuron pool identity and target-muscle connectivity. Cell 123, 477–491 (2005). 10.1016/j.cell.2005.09.009

2 Dasen, J. S., Liu, J. P. & Jessell, T. M. Motor neuron columnar fate imposed by sequential phases of Hox-c activity. Nature 425, 926–933 (2003). 10.1038/nature02051

3 Philippidou, P. & Dasen, J. S. Hox genes: choreographers in neural development, architects of circuit organization. Neuron 80, 12–34 (2013). 10.1016/j.neuron.2013.09.020

4 Jung, H. et al. Evolving Hox activity profiles govern diversity in locomotor systems. Dev Cell 29, 171–187 (2014). 10.1016/j.devcel.2014.03.008

5 Miller, A. & Dasen, J. S. Establishing and maintaining Hox profiles during spinal cord development. Semin Cell Dev Biol 152-153, 44–57 (2024). 10.1016/j.semcdb.2023.03.014

6 Sances, S. et al. Modeling ALS with motor neurons derived from human induced pluripotent stem cells. Nat Neurosci 19, 542–553 (2016). 10.1038/nn.4273

7 Thiry, L., Hamel, R., Pluchino, S., Durcan, T. & Stifani, S. Characterization of Human iPSC-derived Spinal Motor Neurons by Single-cell RNA Sequencing. Neuroscience 450, 57–70 (2020). 10.1016/j.neuroscience.2020.04.041

8 Ho, R. et al. Cross-Comparison of Human iPSC Motor Neuron Models of Familial and Sporadic ALS Reveals Early and Convergent Transcriptomic Disease Signatures. Cell Syst 12, 159–175 e159 (2021). 10.1016/j.cels.2020.10.010

9 Maury, Y. et al. Combinatorial analysis of developmental cues efficiently converts human pluripotent stem cells into multiple neuronal subtypes. Nature Biotechnology 33, 89–U270 (2015). 10.1038/nbt.3049

10 Workman, M. J. et al. Large-scale differentiation of iPSC-derived motor neurons from ALS and control subjects. Neuron 111, 1191–1204 e1195 (2023). 10.1016/j.neuron.2023.01.010

11 Liu, J. P., Laufer, E. & Jessell, T. M. Assigning the positional identity of spinal motor neurons: rostrocaudal patterning of Hox-c expression by FGFs, Gdf11, and retinoids. Neuron 32, 997–1012 (2001). 10.1016/s0896-6273(01)00544-x

12 Lippmann, E. S. et al. Deterministic Patterning in Human Pluripotent Stem Cell-Derived Neuroectoderm. Stem Cell Reports 4, 632–644 (2015). 10.1016/j.stemcr.2015.02.018

13 Bok, J. et al. A controllable human spinal cord model with full dorsoventral patterning. Nat Commun 17 (2026). 10.1038/s41467-026-71162-z

14 Cooper, F. et al. Notch signalling influences cell fate decisions and HOX gene induction in axial progenitors. Development 151 (2024). 10.1242/dev.202098

15 Diez Del Corral, R. & Morales, A. V. The Multiple Roles of FGF Signaling in the Developing Spinal Cord. Front Cell Dev Biol 5, 58 (2017). 10.3389/fcell.2017.00058

16 Henrique, D., Abranches, E., Verrier, L. & Storey, K. G. Neuromesodermal progenitors and the making of the spinal cord. Development 142, 2864–2875 (2015). 10.1242/dev.119768

17 Gouti, M. et al. In vitro generation of neuromesodermal progenitors reveals distinct roles for wnt signalling in the specification of spinal cord and paraxial mesoderm identity. PLoS Biol 12, e1001937 (2014). 10.1371/journal.pbio.1001937

18 Mouilleau, V. et al. Dynamic extrinsic pacing of the HOX clock in human axial progenitors controls motor neuron subtype specification. Development 148 (2021). 10.1242/dev.194514

19 Baxi, E. G. et al. Answer ALS, a large-scale resource for sporadic and familial ALS combining clinical and multi-omics data from induced pluripotent cell lines. Nat Neurosci 25, 226–237 (2022). 10.1038/s41593-021-01006-0

20 Loh, L. J. et al. Generation of spinal cord organoids from human induced pluripotent stem cells caudalised to a lumbar fate. Sci Rep (2026). 10.1038/s41598-026-45679-8

21 Lim, G. S. et al. Microhexagon gradient array directs spatial diversification of spinal motor neurons. Theranostics 9, 311–323 (2019). 10.7150/thno.29755

22 Patani, R. et al. Retinoid-independent motor neurogenesis from human embryonic stem cells reveals a medial columnar ground state. Nat Commun 2, 214 (2011). 10.1038/ncomms1216

23 Lin, H. C. et al. Human neuron subtype programming via single-cell transcriptome-coupled patterning screens. Science 389, eadn6121 (2025). 10.1126/science.adn6121

24 Xu, H. J. et al. Generation of functional posterior spinal motor neurons from hPSCs-derived human spinal cord neural progenitor cells. Cell Regen 12, 15 (2023). 10.1186/s13619-023-00159-6

25 Zhu, Y. et al. Engineered thoracic spinal cord organoids for transplantation after spinal cord injury. Nat Biomed Eng (2025). 10.1038/s41551-025-01549-8

26 Thiry, L., Sirois, J., Durcan, T. M. & Stifani, S. Generation of human iPSC-derived phrenic-like motor neurons to model respiratory motor neuron degeneration in ALS. Commun Biol 7, 238 (2024). 10.1038/s42003-024-05925-z

27 Caldarelli, P. et al. Self-organized tissue mechanics underlie embryonic regulation. Nature 633, 887–894 (2024). 10.1038/s41586-024-07934-8

28 Abdel Fattah, A. R., et al. Actuation enhances patterning in human neural tube organoids. Nat Commun 12, 3192 (2021). 10.1038/s41467-021-22952-0

29 Sareen, D. et al. Human induced pluripotent stem cells are a novel source of neural progenitor cells (iNPCs) that migrate and integrate in the rodent spinal cord. J Comp Neurol 522, 2707–2728 (2014). 10.1002/cne.23578

30 Shelley, B. C., Gowing, G. & Svendsen, C. N. A cGMP-applicable expansion method for aggregates of human neural stem and progenitor cells derived from pluripotent stem cells or fetal brain tissue. J Vis Exp (2014). 10.3791/51219

31 Svendsen, C. N. et al. A new method for the rapid and long term growth of human neural precursor cells. J Neurosci Methods 85, 141–152 (1998). 10.1016/s0165-0270(98)00126-5

32 Mendelsohn, A. I., Dasen, J. S. & Jessell, T. M. Divergent Hox Coding and Evasion of Retinoid Signaling Specifies Motor Neurons Innervating Digit Muscles. Neuron 93, 792–805 e794 (2017). 10.1016/j.neuron.2017.01.017

33 Adams, K. L., Rousso, D. L., Umbach, J. A. & Novitch, B. G. Foxp1-mediated programming of limb-innervating motor neurons from mouse and human embryonic stem cells. Nature Communications 6 (2015). ARTN 6778 10.1038/ncomms7778

34 Rousso, D. L., Gaber, Z. B., Wellik, D., Morrisey, E. E. & Novitch, B. G. Coordinated actions of the forkhead protein Foxp1 and Hox proteins in the columnar organization of spinal motor neurons. Neuron 59, 226–240 (2008). 10.1016/j.neuron.2008.06.025

35 Palmesino, E. et al. Foxp1 and lhx1 coordinate motor neuron migration with axon trajectory choice by gating Reelin signalling. PLoS Biol 8, e1000446 (2010). 10.1371/journal.pbio.1000446

36 Pattyn, A., Hirsch, M., Goridis, C. & Brunet, J. F. Control of hindbrain motor neuron differentiation by the homeobox gene Phox2b. Development 127, 1349–1358 (2000). 10.1242/dev.127.7.1349

37 Lawrence, J. E. G. et al. HOX gene expression in the developing human spine. Nat Commun 15, 10023 (2024). 10.1038/s41467-024-54187-0

38 Li, E. et al. CRISPRi-based screens in iAssembloids to elucidate neuron-glia interactions. Neuron 113, 701–718 e708 (2025). 10.1016/j.neuron.2024.12.016

39 Frega, M., Tedesco, M., Massobrio, P., Pesce, M. & Martinoia, S. Network dynamics of 3D engineered neuronal cultures: a new experimental model for in-vitro electrophysiology. Sci Rep 4, 5489 (2014). 10.1038/srep05489

40 Horberg, C. J., Englund Johansson, U., Johansson, F. & O’Carroll, D. Spontaneous Cell Cluster Formation in Human iPSC-Derived Neuronal Spheroid Networks Influences Network Activity. eNeuro 9 (2022). 10.1523/ENEURO.0143-22.2022

41 Andersen, J. et al. Generation of Functional Human 3D Cortico-Motor Assembloids. Cell 183, 1913–1929 e1926 (2020). 10.1016/j.cell.2020.11.017

42 Pereira, J. D. et al. Human sensorimotor organoids derived from healthy and amyotrophic lateral sclerosis stem cells form neuromuscular junctions. Nat Commun 12, 4744 (2021). 10.1038/s41467-021-24776-4

43 Galluzzi, G., Ruocco, G., Fornetti, E. & Genovese, I. Modeling ALS in a dish: how organoids are transforming research. Front Med (Lausanne) 13, 1792336 (2026). 10.3389/fmed.2026.1792336

44 Eigenhuis, K. N., Ferrer, R. M. & Pasterkamp, R. J. Neural Organoid Models as a Platform for Studying Disease Mechanisms in Amyotrophic Lateral Sclerosis. J Neurochem 170, e70513 (2026). 10.1111/jnc.70513

45 Takazawa, T. et al. Maturation of spinal motor neurons derived from human embryonic stem cells. PLoS One 7, e40154 (2012). 10.1371/journal.pone.0040154

46 Taga, A. et al. Role of Human-Induced Pluripotent Stem Cell-Derived Spinal Cord Astrocytes in the Functional Maturation of Motor Neurons in a Multielectrode Array System. Stem Cells Transl Med 8, 1272–1285 (2019). 10.1002/sctm.19-0147

47 Qu, Q., et al. High-efficiency motor neuron differentiation from human pluripotent stem cells and the function of Islet-1. Nat Commun 5, 3449 (2014). 10.1038/ncomms4449

48 Tadros, M. A., Fuglevand, A. J., Brichta, A. M. & Callister, R. J. Intrinsic excitability differs between murine hypoglossal and spinal motoneurons. J Neurophysiol 115, 2672–2680 (2016). 10.1152/jn.01114.2015

49 Smith, C. C. & Brownstone, R. M. Spinal motoneuron firing properties mature from rostral to caudal during postnatal development of the mouse. J Physiol 598, 5467–5485 (2020). 10.1113/JP280274

50 Parodi, G., Brofiga, M., Pastore, V. P., Chiappalone, M. & Martinoia, S. Deepening the role of excitation/inhibition balance in human iPSCs-derived neuronal networks coupled to MEAs during long-term development. J Neural Eng 20 (2023). 10.1088/1741-2552/acf78b

51 Buccino, A. P. et al. SpikeInterface, a unified framework for spike sorting. Elife 9 (2020). 10.7554/eLife.61834

52 Garcia, S. et al. Opening the black box: a modular approach to spike sorting. bioRxiv (2026). 10.64898/2026.01.23.701239

53 Hornauer, P. et al. DeePhys: A machine learning-assisted platform for electrophysiological phenotyping of human neuronal networks. Stem Cell Reports 19, 285–298 (2024). 10.1016/j.stemcr.2023.12.008

