## Supplemental Figures 1-8 and legends for "Generation of human iPSC-derived motor neurons with defined regional identities across the rostrocaudal axis"

**Supplemental Figure 1**

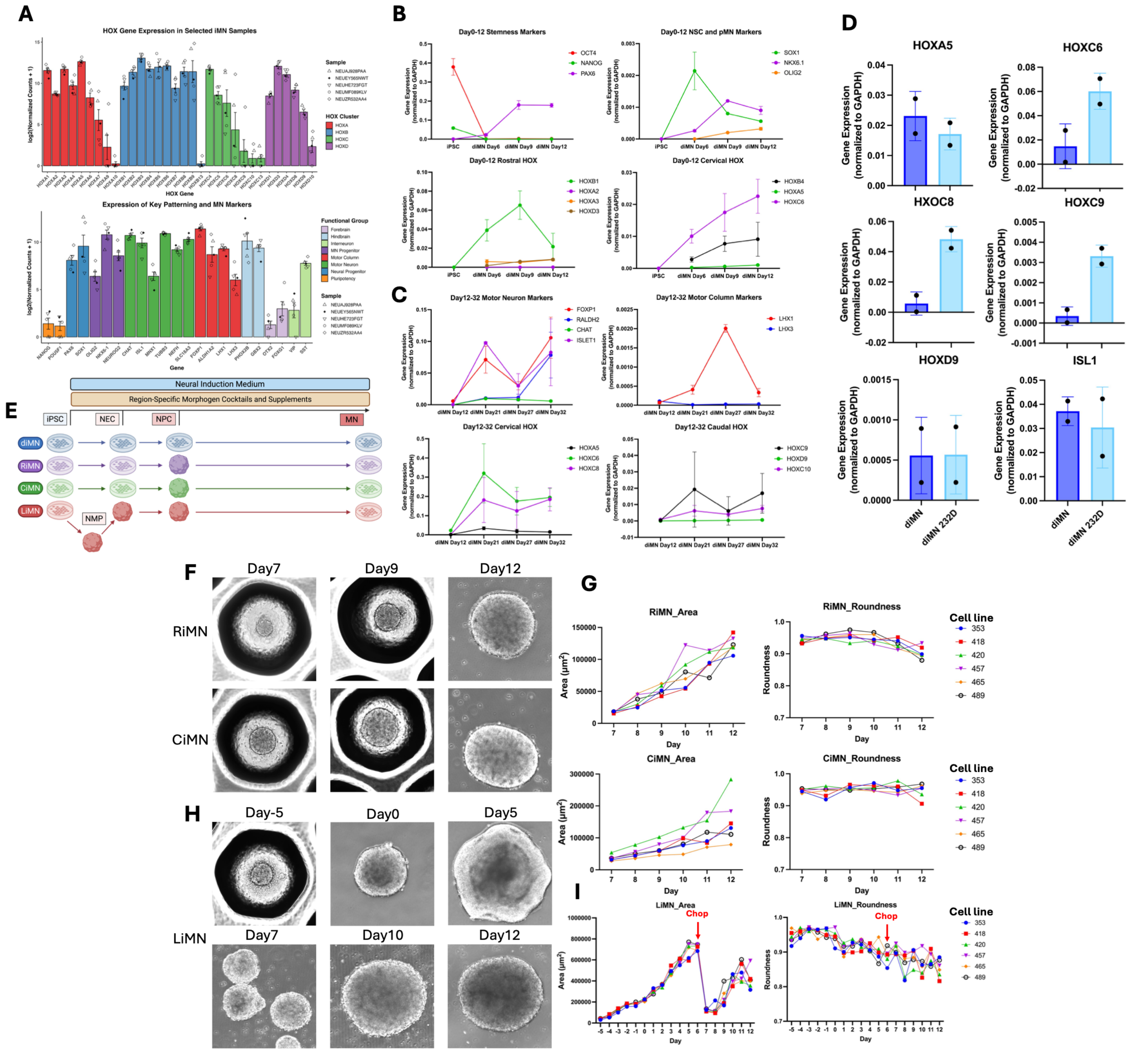


**Supplemental Figure 1. Characterization of conventional iMN differentiation and transient spheroid culture during regional patterning. Related to Figure 1. (A)** Expression of HOX genes (top) and selected regional, neural, and motor neuron markers (bottom) in five selected diMN samples from the Answer ALS dataset. Expression is displayed as $\log_2$. Colors indicate HOX clusters or marker categories, and symbols identify individual samples. Bars show mean ± SD. **(B)** RT-qPCR time courses of pluripotency, neural progenitor, and rostrocaudal patterning markers during conventional diMN differentiation, showing undifferentiated iPSCs and differentiation days 6, 9, and 12. **(C)** RT-qPCR time courses of motor neuron, motor column, and cervical and caudal HOX markers during diMN maturation at days 12, 21, 27, and 32. Data in **(B)** and **(C)** show mean ± SD across three independent iPSC lines. **(D)** RT-qPCR comparison of selected HOX genes and *ISL1* at day 32 between conventional adherent diMNs and diMNs generated using a transient 3D-to-2D culture format (diMN 232D). Each point represents one independent iPSC line (n = 2 per condition), and bars show mean ± SD. Expression in **(B–D)** was normalized to *GAPDH* using the $2^{-\Delta Ct}$ method. **(E)** Schematic of the culture formats used to generate diMNs, RiMNs, CiMNs, and LiMNs. Regional protocols incorporate a transient spheroid phase followed by adherent maturation, with LiMN differentiation proceeding through a neuromesodermal progenitor-like intermediate. NEC, neuroepithelial cell; NPC, neural progenitor cell; NMP, neuromesodermal progenitor; MN, motor neuron. **(F)** Representative brightfield images of RiMN (top) and CiMN (bottom) spheroids at days 7, 9, and 12. **(G)** Projected area and roundness of RiMN and CiMN spheroids from days 7–12. **(H)** Representative brightfield images of LiMN spheroids at days −5, 0, 5, 7, 10, and 12, showing spheroid growth and morphology before and after mechanical division. **(I)** Projected area and roundness of LiMN spheroids from days −5 to 12. "Chop" indicates mechanical division of spheroids. In **(G)** and **(I)**, each colored trajectory represents one of six independent iPSC lines. Each point in **(G)** and **(I)** represents the mean of approximately 10 spheroids per iPSC line at the indicated time point. Source measurements are provided in Table S4.

**Supplemental Figure 2**

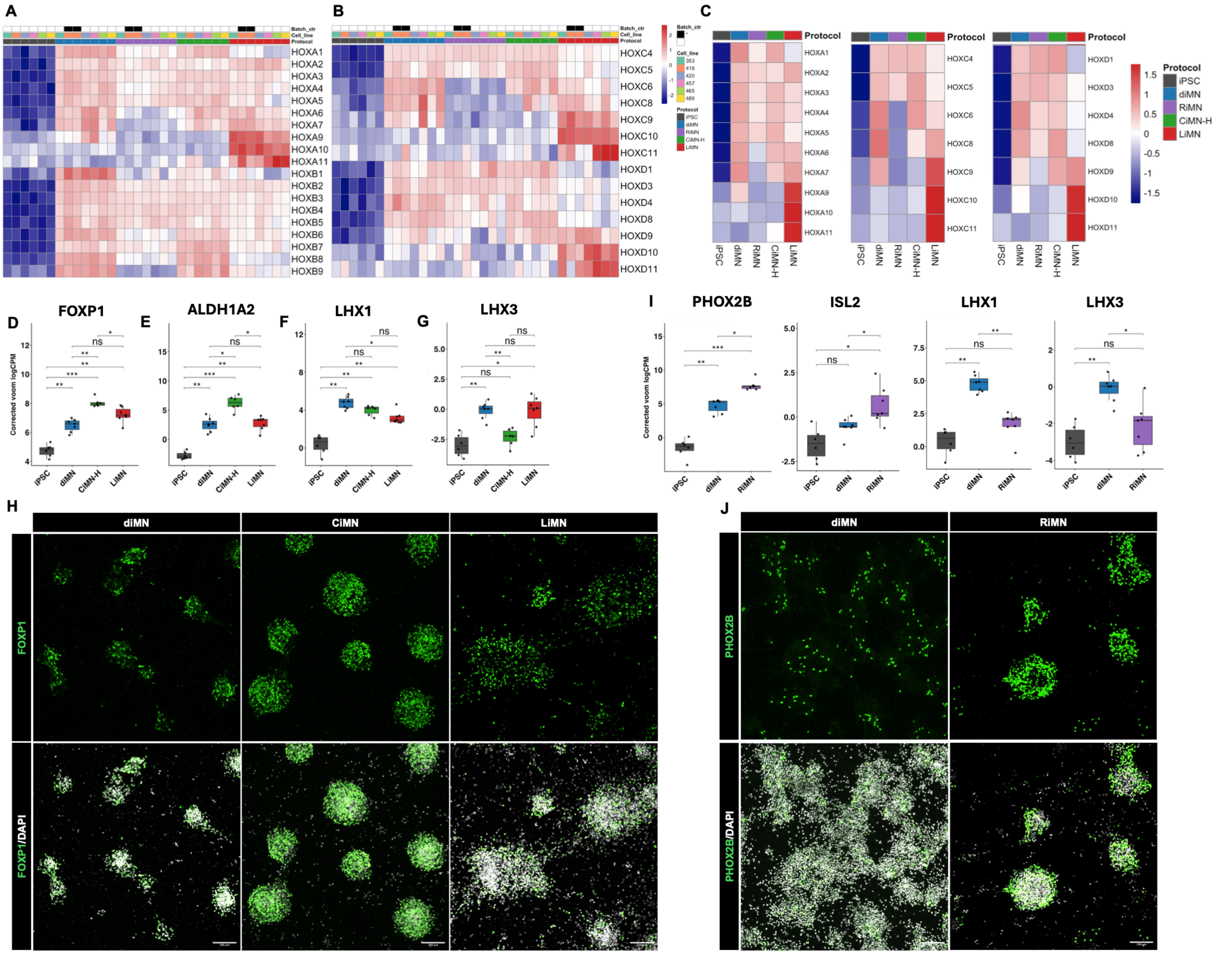


**Supplemental Figure 2. Regional HOX expression and motor neuron subtype-associated features of region-specific iMNs. Related to Figure 2. (A and B)** Heatmaps of HOXA and HOXB **(A)** and HOXC and HOXD **(B)** expression in iPSCs and the four iMN populations at day 32. Columns represent individual samples, with annotations indicating protocol, cell line, and batch-control status. Batch- and cell-line-corrected voom $log_2$ counts per million were standardized as gene-wise z-scores. Genes are ordered by HOX paralog number. **(C)** Heatmaps of protocol-averaged HOXA, HOXC, and HOXD expression, displayed as gene-wise z-scores. **(D–G)** Bulk RNA-seq expression of *FOXP1* **(D)**, *ALDH1A2* **(E)**, *LHX1* **(F)**, and *LHX3* **(G)** in iPSCs and day-32 diMNs, CiMNs, and LiMNs. **(H)** Representative immunofluorescence images of FOXP1 (green) in day-32 diMNs, CiMNs, and LiMNs, with DAPI counterstaining (white). **(I)** Bulk RNA-seq expression of *ISL2*, *PHOX2B*, *LHX1*, and *LHX3* in iPSCs and day-32 diMNs and RiMNs. **(J)** Representative immunofluorescence images of PHOX2B (green) in day-32 diMNs and RiMNs, with DAPI counterstaining (white). In **(D–G)** and **(I)**, each point represents one independent iPSC line. Expression is displayed as batch- and cell-line-corrected voom $log_2$ counts per million. Statistical significance was assessed using two-sided paired t-tests across matched cell lines, with Benjamini–Hochberg correction within each marker-analysis set. *Adjusted $P < 0.05$; **adjusted $P < 0.01$; ***adjusted $P < 0.001$; ns, not significant. Scale bars in **(H)** and **(J)**, 100 μm.

### Supplemental Figure 3

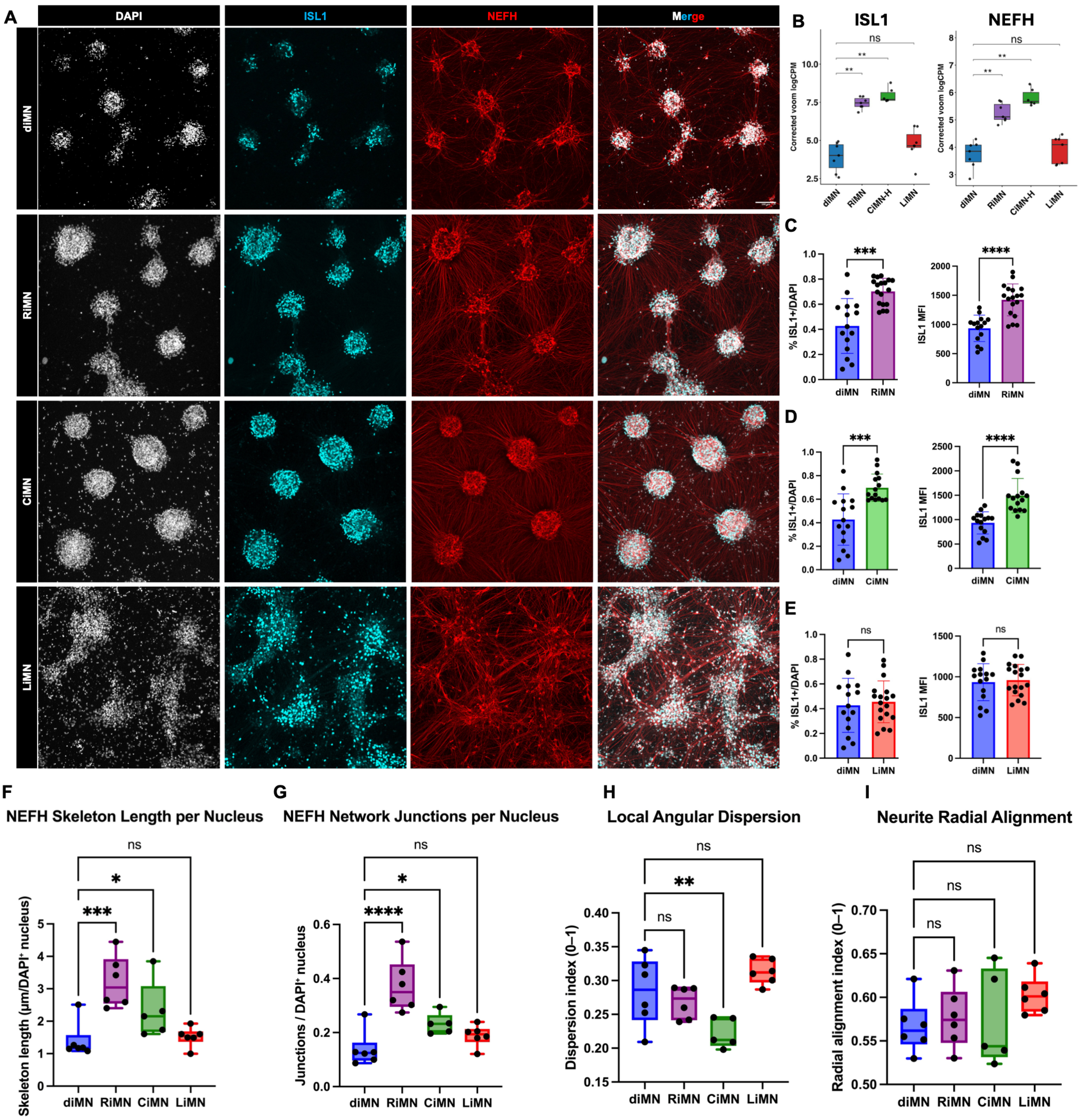


**Supplemental Figure 3. Motor neuron marker expression and neurite-network morphology in region-specific iMNs. Related to Figure 2. (A)** Representative immunofluorescence images of day-32 diMNs, RiMNs, CiMNs, and LiMNs stained for ISL1 (cyan) and NEFH (red), with DAPI counterstaining (white). Merged images are shown in the rightmost column. Scale bars, 100 µm. **(B)** Bulk RNA-seq expression of *ISL1* and *NEFH* at day 32 across the four iMN populations, displayed as batch- and cell-line-corrected voom $\log_2$ counts per million. Each point represents one independent iPSC line. Statistical significance was assessed using two-sided paired t-tests across matched cell lines, with Benjamini–Hochberg correction for multiple comparisons. **(C–E)** Quantification of the fraction of ISL1-positive cells among DAPI-positive nuclei (left) and ISL1 mean fluorescence intensity (right) at day 32, comparing RiMNs **(C)**, CiMNs **(D)**, and LiMNs **(E)** with diMNs. Each point represents one image, and bars show mean ± SD. Statistical significance was assessed using Welch's t-test.

**(F and G)** Quantification of NEFH-positive neurite skeleton length **(F)** and 2D network junctions **(G)** at day 32, normalized to the number of DAPI-positive nuclei. **(H and I)** Quantification of local angular dispersion **(H)** and radial alignment **(I)** of NEFH-positive processes at day 32. Lower angular dispersion indicates greater local alignment, whereas higher radial alignment indicates greater alignment with the radial direction from the identified centers. For **(F–I)**, measurements from three images per cell line and condition were averaged to obtain one value per independent iPSC line (n = 5 for CiMNs; n = 6 for diMNs, RiMNs, and LiMNs). Boxes show the median and interquartile range, with whiskers indicating the minimum and maximum values. Statistical significance was assessed using one-way ANOVA followed by Dunnett's multiple-comparisons test, comparing each regional iMN population with the diMN control. Source measurements are provided in Table S5. $^{*}P < 0.05$; $^{**}P < 0.01$; $^{***}P < 0.001$; $^{****}P < 0.0001$; ns, not significant. Significance annotations in **(B)** indicate adjusted P-values.

### Supplemental Figure 4

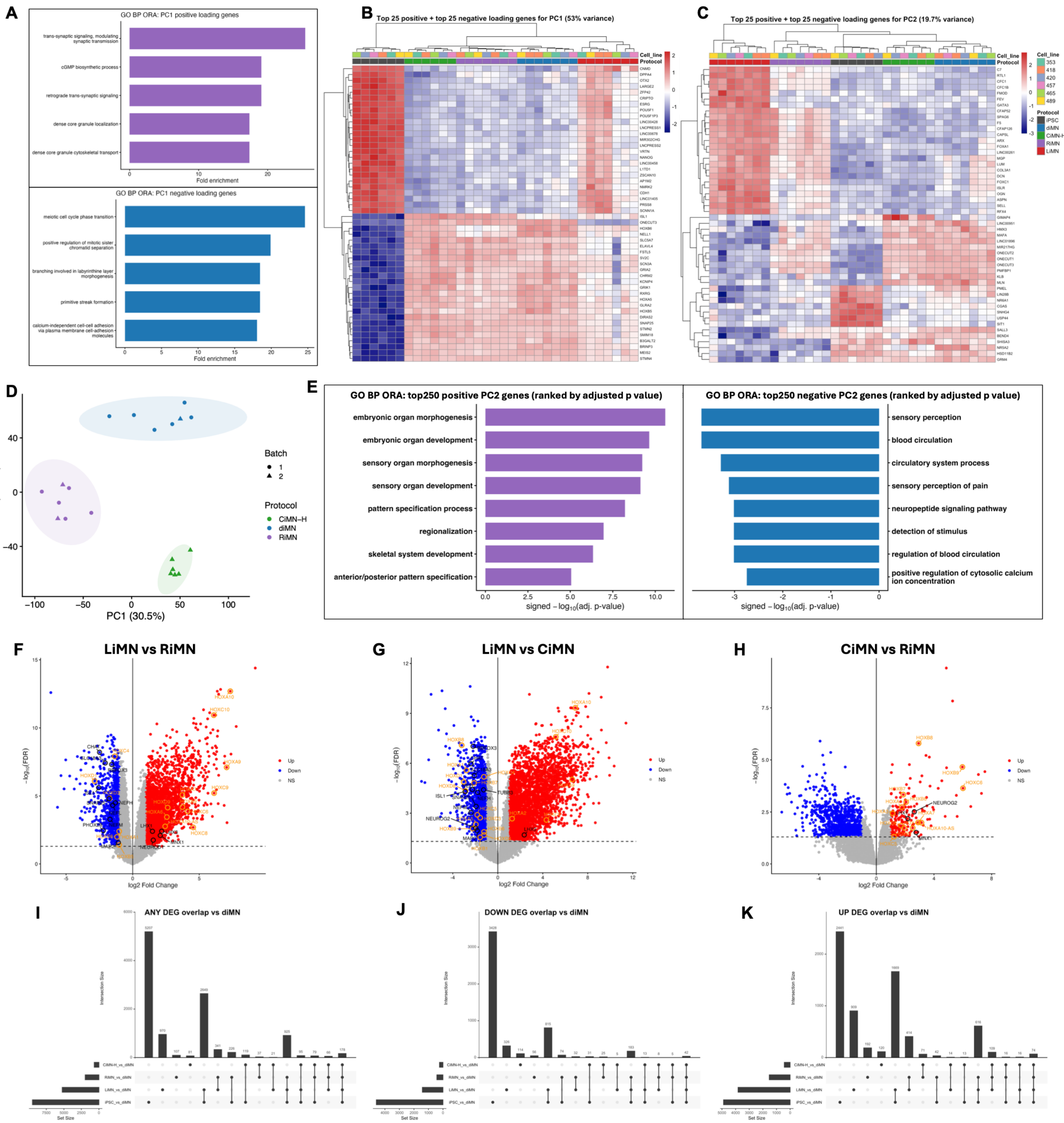

**Supplemental Figure 4. Principal component loading analyses and differential gene expression across iMN populations. (A)** Gene Ontology (GO) Biological Process over-representation analysis of the 250 genes with the strongest positive (top) and negative (bottom) PC1 loadings from the PCA in Figure 3A. The five highest-ranking terms by fold enrichment are displayed for each direction. **(B and C)** Heatmaps of the 25 genes with the strongest positive and 25 genes with the strongest negative loadings for PC1 **(B)** and PC2 **(C)**. Rows represent genes, and columns represent individual samples. Colors indicate gene-wise z-scores of batch- and cell-line-corrected voom $log_2$ counts per million. Column annotations indicate cell line and differentiation protocol. **(D)** PCA refitted using only day-32 diMN, RiMN, and CiMN samples. Each point represents one sequenced sample. Colors indicate differentiation protocol, and symbols indicate sequencing batch. Shaded ellipses indicate 95% confidence regions.

**(E)** GO Biological Process over-representation analysis of the 250 genes with the strongest positive (left) and negative (right) PC2 loadings from the restricted PCA in **(D)**. The eight highest-ranking terms by adjusted P-value are displayed for each direction. **(F–H)** Volcano plots of differential gene expression in LiMNs relative to RiMNs **(F)**, LiMNs relative to CiMNs **(G)**, and CiMNs relative to RiMNs **(H)**. Each point represents one gene, plotted by $\log_2$ fold change and $-\log_{10}$ false discovery rate (FDR). Red and blue indicate upregulated and downregulated genes, respectively, defined by FDR < 0.05 and absolute $\log_2$ fold change > 1. Gray indicates genes not meeting both thresholds. **(I–K)** UpSet plots showing overlaps among all differentially expressed genes **(I)**, downregulated genes **(J)**, and upregulated genes **(K)** in iPSCs, RiMNs, CiMNs, and LiMNs relative to diMNs, using FDR < 0.05 and absolute $\log_2$ fold change > 1. Horizontal bars indicate gene-set sizes, vertical bars indicate intersection sizes, and connected dots identify the sets contributing to each intersection.

**Supplemental Figure 5**

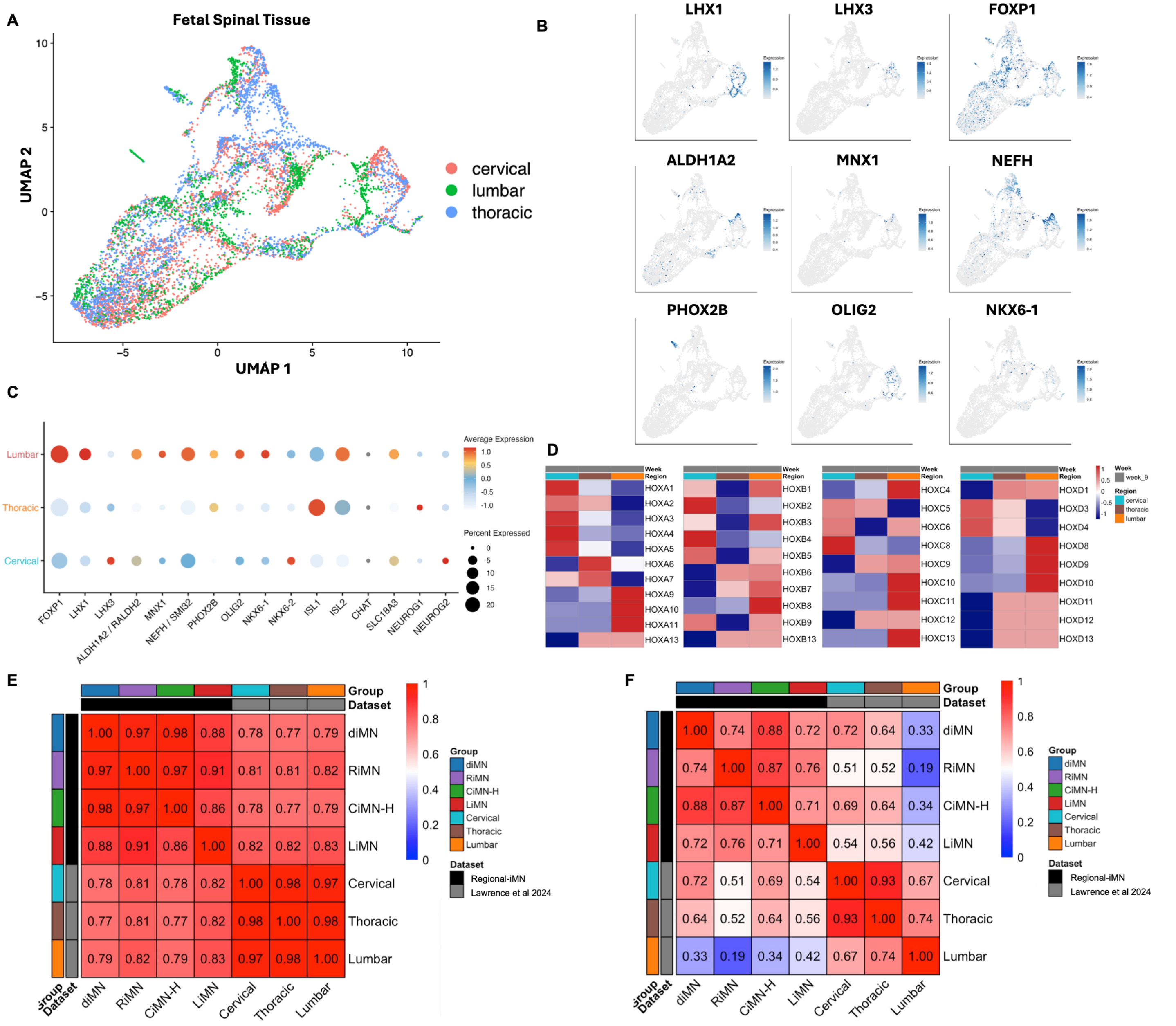


**Supplemental Figure 5. Characterization of human fetal spinal cord reference profiles and comparison with regional iMNs. (A)** UMAP of week-9 human fetal spinal cord spatial transcriptomic profiles, colored by cervical, thoracic, and lumbar anatomical regions. **(B)** UMAP feature plots showing normalized expression of selected motor neuron, progenitor, and subtype-associated markers. **(C)** Dot plot of selected marker-gene expression across cervical, thoracic, and lumbar neuron/glia-enriched spatial spots. Dot size indicates the percentage of spots expressing each gene, and color indicates scaled average expression. **(D)** Heatmaps of HOX gene expression across week-9 fetal cervical, thoracic, and lumbar regional pseudobulk profiles. Colors indicate gene-wise z-scores. **(E and F)** Spearman correlation heatmaps comparing day-32 diMNs, RiMNs, CiMNs, and LiMNs with week-9 fetal cervical, thoracic, and lumbar reference populations using all shared genes **(E)** or shared HOX genes together with motor neuron and progenitor markers **(F)**. Colors and values indicate Spearman correlation coefficients.

### Supplemental Figure 6

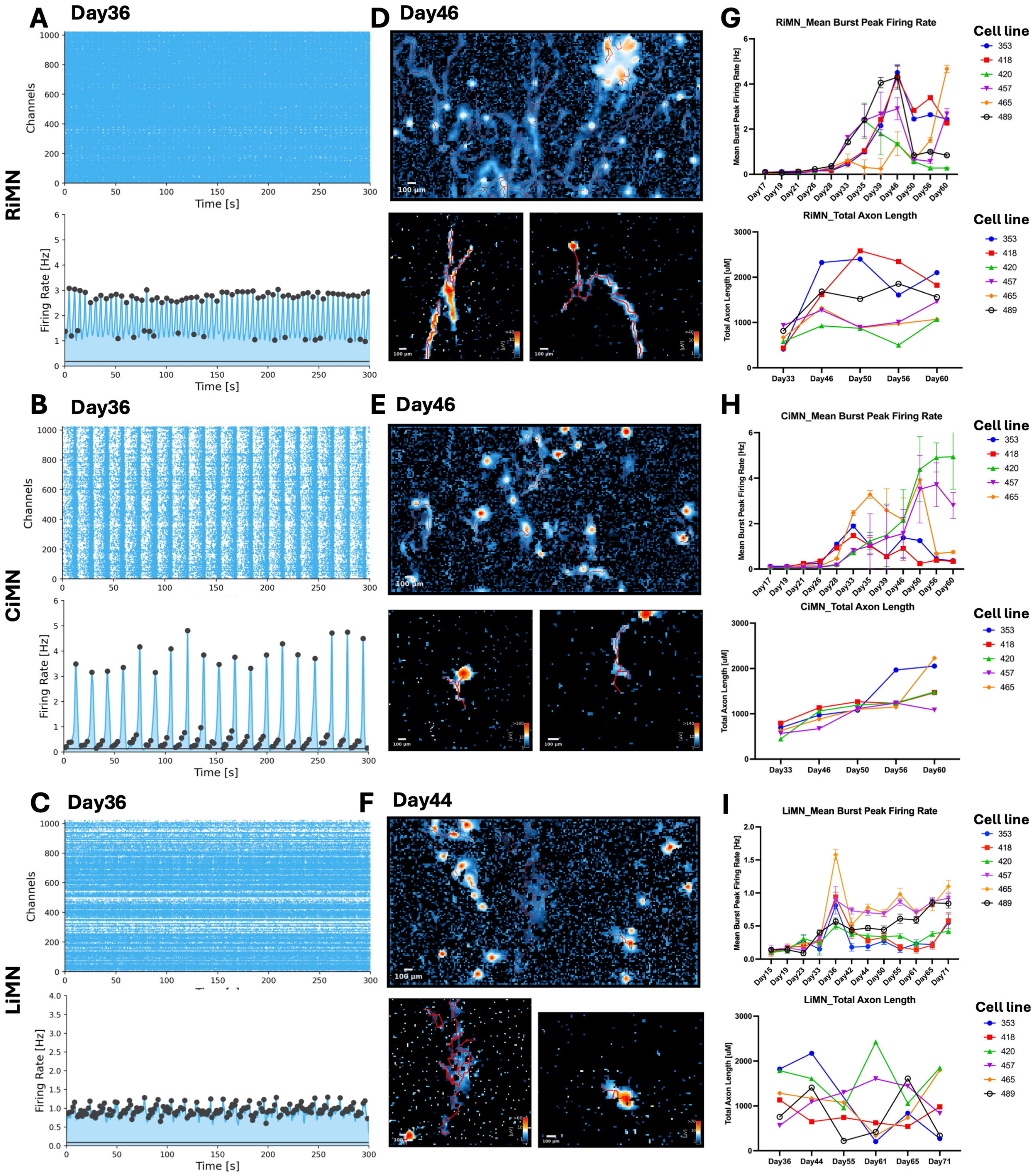


**Supplemental Figure 6. Additional HD-MEA recordings and longitudinal functional measurements in regional iMNs. Related to Figure 5. (A–C)** Representative spike raster plots (top) and corresponding network firing-rate traces (bottom) from additional RiMN (A), CiMN (B), and LiMN (C) cultures at day 36. Raster plots show spike events across recording channels over 300 s. **(D–F)** Representative axon-tracking maps from RiMNs at day 46 (D), CiMNs at day 46 (E), and LiMNs at day 44 (F). Upper panels show full-field activity maps, and lower panels show examples of individual neuronal electrical footprints with tracked axonal trajectories. Scale bars, 100 µm. **(G–I)** Longitudinal measurements of mean burst peak firing rate (top) and total axon length (bottom) in RiMNs (G), CiMNs (H), and LiMNs (I). Each colored trajectory represents one independent iPSC line (n = 5 for CiMNs; n = 6 for RiMNs and LiMNs). Error bars, where shown, indicate SD, and differentiation days are indicated on the x-axis.

### Supplemental Figure 7

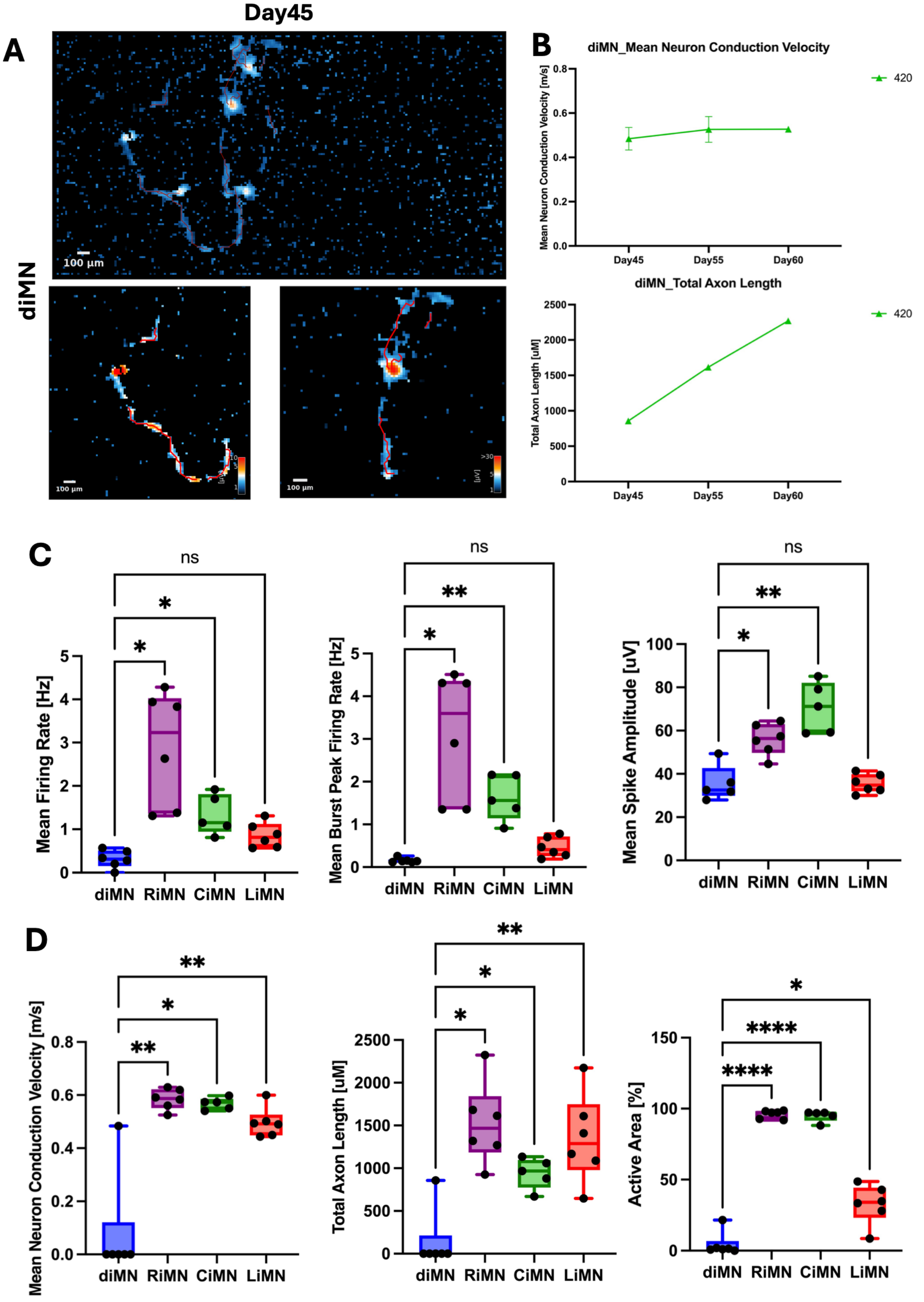


**Supplemental Figure 7. Axonal activity in conventional diMNs and functional comparisons across iMN populations. Related to Figure 5. (A)** Representative axon-tracking maps from diMNs at day 45. The upper panel shows a full-field activity map, and lower panels show examples of individual neuronal electrical footprints with tracked axonal trajectories. Scale bars, 100 µm. **(B)** Longitudinal measurements of mean neuronal conduction velocity (top) and total axon length (bottom) in diMNs derived from cell line 420 at days 45, 55, and 60. Error bars indicate SD. **(C)** Comparison of mean firing rate (left), mean burst peak firing rate (middle), and mean spike amplitude (right) across diMNs, RiMNs, CiMNs, and LiMNs at approximately day 45. **(D)**

Comparison of mean neuronal conduction velocity (left), total axon length (middle), and electrically active area (right) across the four iMN populations at approximately day 45. In **(C)** and **(D)**, each point represents one independent iPSC line (n = 5 for CiMNs; n = 6 for diMNs, RiMNs, and LiMNs). Boxes show the median and interquartile range, with whiskers indicating the minimum and maximum values. Statistical significance was assessed using mixed-effects analysis with cell line as the matching factor and Geisser–Greenhouse correction, followed by Dunnett's multiple-comparisons test comparing each regional iMN population with diMNs. *$P < 0.05$; **$P < 0.01$; ***$P < 0.001$; ****$P < 0.0001$; ns, not significant.

**Supplemental Figure 8**

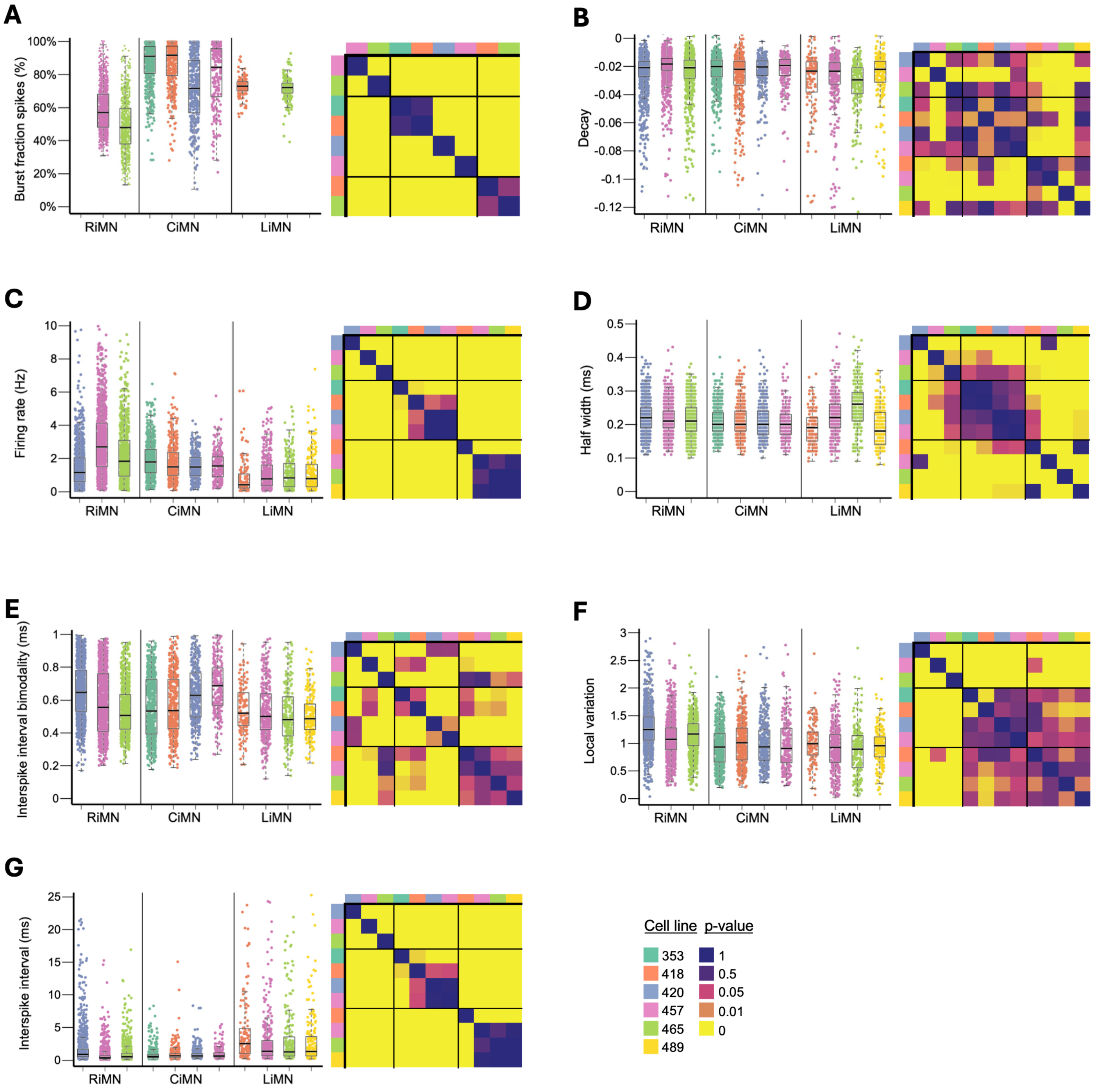


**Supplemental Figure 8. Single-neuron electrophysiological feature distributions across regional iMN cultures and cell lines.** Spike-sorted HD-MEA recordings acquired at day 45 were analyzed for **(A)** burst fraction of spikes, **(B)** waveform decay, **(C)** firing rate, **(D)** spike half-width, **(E)** interspike interval bimodality, **(F)** local variation, and **(G)** mean interspike interval. Scatter plots with overlaid boxplots show unit-level feature distributions for individual cell lines within each regional protocol. Each point represents one spike-sorted unit, and colors indicate donor cell line. Burst fraction values were unavailable for RiMN cell line 420 and LiMN cell lines 457 and 489; these cultures were therefore omitted from panel A. Adjacent heatmaps display Benjamini–Hochberg-adjusted P-values from pairwise Kolmogorov–Smirnov tests comparing feature distributions between wells. Colored annotation bars identify donor cell lines, and black boundaries delineate regional protocol groups. Missing feature values were excluded from the corresponding analyses. Unit-level data are provided in Table S13.
